# Postnatal maternal care moderates the effects of prenatal bisphenol exposure on offspring neurodevelopmental, behavioral, and transcriptomic outcomes

**DOI:** 10.1101/2023.09.19.558481

**Authors:** Samantha C. Lauby, Hannah E. Lapp, Melissa Salazar, Sofiia Semyrenko, Danyal Chauhan, Amy E. Margolis, Frances A. Champagne

## Abstract

Bisphenols (BPs), including BPA and “BPA-free” structural analogs, are commonly used plasticizers that are present in many plastics and are known endocrine disrupting chemicals. Prenatal exposure to BPA has been associated with negative neurodevelopmental and behavioral outcomes in children and rodent models. Prenatal BPA exposure has also been shown to impair postnatal maternal care provisioning, which can also affect offspring neurodevelopment and behavior. However, there is limited knowledge regarding the biological effects of prenatal exposure to bisphenols other than BPA and the interplay between prenatal BP exposure and postnatal maternal care on adult behavior. The purpose of the current study was to determine the interactive impact of prenatal BP exposure and postnatal maternal care on neurodevelopment and behavior. Our findings suggest that the effects of prenatal BP exposure on eye-opening, adult attentional set shifting and anxiety-like behavior in the open field are dependent on maternal care in the first five days of life. Interestingly, maternal care might also attenuate the effects of prenatal BP exposure on eye opening and adult attentional set shifting. Finally, transcriptomic profiles in male and female medial prefrontal cortex and amygdala suggest that the interactive effects of prenatal BP exposure and postnatal maternal care converge on estrogen receptor signaling and are involved in biological processes related to gene expression and protein translation and synthesis. Overall, these findings indicate that postnatal maternal care plays a critical role in the expression of the effects of prenatal BP exposure on neurodevelopment and adult behavior. Understanding the underlying biological mechanisms involved might allow us to identify potential avenues to mitigate the adverse effects of prenatal BP exposure and improve health and well-being in human populations.

## 1. Introduction

Environmental exposures occurring during early-life, including the prenatal and early postnatal period, can have a profound influence on neurodevelopmental trajectories and behavior that persist into adulthood. Though experimental studies of environmental toxicants have typically focused on exposure to a single chemical, there is increasing appreciation of the need to consider the exposome, which represents the collective impact of environmental exposures and includes both risk and protective factors (Appleton et al., 2016; Tamiz et al., 2022). Evidence for the complex interactions between exposures in predicting neurodevelopmental outcomes is emerging and suggests significant interplay between early life environmental and social experiences (Appleton et al., 2016). Understanding the nature of this interplay will be critical to the mechanistic understanding of risk and resilience.

Endocrine disrupting chemicals, such as bisphenols (BPs), are pervasive in everyday life and may be particularly disruptive to neurodevelopment. BPs such as BPA and other “BPA-free” structural analogs, including BPF and BPS, have been shown to act as endocrine disruptors that primarily affect estrogen receptor signaling. BPs are commonly used plasticizers that are present in many plastics, such as food storage containers and reusable water bottles, as well as the lining of food cans. BP exposure via oral consumption is ubiquitous in human populations with inter-individual variation in the levels of exposure (Huang et al., 2017; Lehmler et al., 2018; Lorber et al., 2015). Prenatal exposure to BPs has been associated with negative neurodevelopmental and behavior outcomes for children in human epidemiological studies (Bornehag et al., 2021; Braun et al., 2009, 2017; Perera et al., 2012, 2016; Rochester et al., 2018, 2018; Roen et al., 2015) as well as in rodent models (Arambula et al., 2018; Castro et al., 2015; Kubo et al., 2003; Kundakovic et al., 2013; Ohtani et al., 2017; Wolstenholme et al., 2011). Studies often report a nonmonotonic dose response curve between prenatal BP exposure levels and neurodevelopmental outcomes, such that lower exposure levels are linked to worse outcomes (Vandenberg, 2014; Vandenberg et al., 2013, 2019). In addition, there are often sex-specific effects of prenatal BP exposure on behavior outcomes, possibly due to BP-associated disruption to estrogen-dependent sexual differentiation of the brain occurring early in life (Arambula et al., 2018; Kubo et al., 2003; Kundakovic et al., 2013). Currently, most studies on the effects of prenatal BP exposure are focused on BPA with less known regarding the effects of prenatal exposure to other bisphenols and their mixtures.

Studies using animal models and human placental tissue have demonstrated that BPs can cross the placenta to exert effects directly on the fetal brain (Balakrishnan et al., 2010; Takahashi & Oishi, 2000). In addition, maternal BPs may disrupt estrogen-dependent changes in the maternal brain which impairs postnatal maternal care (Catanese & Vandenberg, 2017; Kundakovic et al., 2013; Seta et al., 2005). Postnatal maternal care, particularly within the first few days of life, has been associated with numerous neurodevelopmental and behavior outcomes in rodent offspring (Beery & Francis, 2011; Champagne et al., 2003; Curley & Champagne, 2016). Studies with longitudinal human birth cohorts suggest similar effects (Hane et al., 2010; Narvaez et al., 2013; Wade et al., 2022). Reduced postnatal maternal care via artificial rearing paradigms is often associated with negative neurodevelopmental and behavior outcomes (Fleming et al., 2002; Lovic et al., 2006; Lovic & Fleming, 2004); therefore, low maternal care could compound the effects of prenatal BP exposure.

The potential modulating influence of postnatal maternal care on prenatal BP exposure effects remains understudied. Previous work from our lab has demonstrated prenatal BPA-induces dose-dependent changes in postnatal maternal care and adult offspring anxiety-like behavior and social behavior in mice (Kundakovic et al., 2013). Postnatal maternal care further altered DNA methylation levels in the estrogen receptor alpha gene as well as the probability of an individual to become socially dominant (Kundakovic et al., 2013). In addition, previous work from our lab has reported that both prenatal exposure to a BP mixture (BPA, BPF, BPS) and postnatal maternal licking/grooming can alter gene expression in the brains of weanling rats (Lapp et al., 2022). Prenatal BP exposure had the strongest effects on the amygdala and postnatal maternal care had the strongest effects on the hypothalamus, suggesting unique exposure-associated effects that may converge to predict neurobehavioral outcomes (Lapp et al., 2022). However, this preliminary work did not explore the interplay between prenatal BP exposure and postnatal maternal care.

The purpose of the current study was to determine the interactive impact of prenatal BP exposure and postnatal maternal care on neurodevelopment and behavior. We exposed pregnant rats to one of two different concentrations of a BP mixture (BPA, BPF, BPS) or BPA only through oral administration and measured home-cage postnatal maternal care. We assessed eye-opening as a neurodevelopmental milestone and measured adult attention (attentional set-shifting performance) and anxiety-like behavior (elevated plus maze and open field). We also examined transcriptomic changes in the developing medial prefrontal cortex and amygdala. These brain regions are strongly associated with attentional set shifting performance (Floresco et al., 2008; Ghods-Sharifi et al., 2008) and anxiety-like behavior (Buijs & Van Eden, 2000), respectively. Previous work has shown that either prenatal BPA exposure or postnatal maternal care can affect eye-opening time (Ellenbroek et al., 2005; Uriarte et al., 2007), adult attentional set shifting performance (Lauby et al., 2019; Lovic & Fleming, 2004; McLean et al., 2010) and anxiety-like behavior (Caldji et al., 2000; Francis & Meaney, 1999; Kundakovic et al., 2013; Ohtani et al., 2017; Wiersielis et al., 2020). Therefore, we hypothesized that interactions between prenatal BP exposure and postnatal maternal care would influence these phenotypes and alter the transcriptome in the developing medial prefrontal cortex and amygdala.

## 2. Materials and Methods

### 2.1 Husbandry and Breeding

All animal procedures were approved by the Institutional Animal Care and Use Committee at the University of Texas at Austin and conformed to the guidelines of the American Association for Laboratory Animal Science. Seven-week-old female (n = 108) and nine-week-old male (n = 26) Long-Evans rats were obtained from Charles River Laboratories and acclimated to the animal facility for two weeks. Rats were housed in same-sex pairs on a 12:12 hour inverse light-dark cycle (lights-off at 10:00 during standard time, 11:00 during daylight savings time) with *ad libitum* access to standard chow diet (#5LL2, Lab Diet) and water. All rats were provided glass water bottles, polysulfone cages, and aspen wood shavings as bedding to limit additional external exposure to bisphenols and other xenoestrogens.

During the mating period (starting at 12:00 for 4-6 days), one male and up to two females were paired and observed for signs of sexual receptivity. Breeding pairs were left to mate overnight if lordosis (female) and mounting (male) were observed. Females were pair-housed with their cage mate following breeding until a few days before parturition (gestational days 16-19). Day of breeding was considered gestational day (GD) 0. Eleven females did not become pregnant using this procedure and were removed from the study. Breeding occurred in batches of 16 to 22 females over the course of 13 months.

### 2.2 Bisphenol Administration

The prenatal treatment groups and experimental timeline can be found in Figure 1A. The prenatal bisphenol treatments were administered as reported in a previous and separate study (Lapp et al., 2022). Cage-pairs of dams were randomly assigned to receive control Corn Oil (n = 22), 50 μg/kg BPA (n = 30), 50 μg/kg Mixed Bisphenols (BP) (n = 23), or 150 μg/kg Mixed BP (n = 22). Bisphenol stocks were created by dissolving 0.1 g of either bisphenol A (BPA; #B0494, TCI, ≥99 %), bisphenol S (BPS; #A17342, Alpha Aesar, ≥99 %), or bisphenol F (BPF; #A11471, Alfa Aesar, ≥98 %) in 10 mL of corn oil (#405435000, Acros Organics). Stocks were well-mixed prior to creating the working stock of bisphenols that were administered to the dams. Working stocks were created by adding the separate bisphenol stocks into 10 mL corn oil (total volume) to the desired final concentration. Equal parts BPA, BPS, and BPF were used for both Mixed BP treatment groups. The final concentrations were 0.15 μg/μl for the 50 μg/kg BPA and 50 μg/kg Mixed BP groups and 0.45 μg/μl for the 150 μg/kg Mixed BP group.

**Figure 1.**
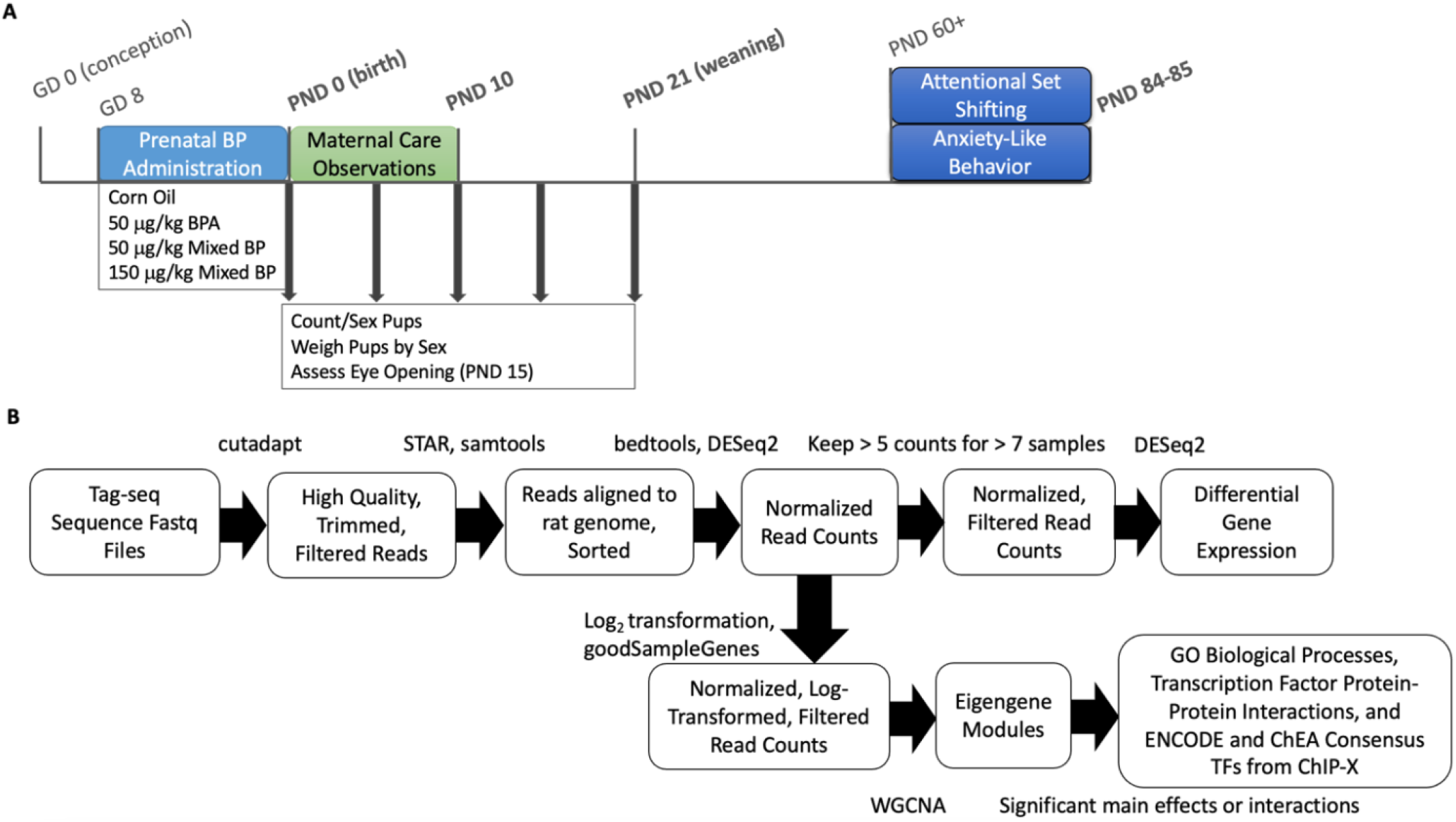
Experimental design, timeline, and Tag-seq analysis pipeline. (A) Timeline for prenatal bisphenol exposure, maternal care observations, and offspring growth, neurodevelopmental and behavioral outcomes. Offspring brains were collected at the bolded ages. (B) Tag-seq analysis pipeline for PND 10 medial prefrontal cortex and amygdala. Fastq files were filtered and trimmed using cutadapt, aligned to the rat genome using STAR, and sorted using samtools. Gene counts were compiled using bedtools and normalized using DEseq2. Differential gene expression analysis was done using DEseq2. Normalized, log-transformed counts were used for weighted gene co-expression network analysis (WGCNA) and the clustered eigengene modules were analyzed. Gene sets from eigengene module with significant main effects or interactions were extracted and analyzed for enriched GO biological processes, Transcription Factor (TF) Protein-Protein Interactions, and ENCODE and ChEA Consensus TFs from ChIP-X.

Following breeding, all dams were provided ¼ of a Nilla wafer with 100 μl of corn oil absorbed into the cookie once a day starting at 8:00 from GD 1 to 7. From GD 8 until parturition (between GD 21-22), dams received ¼ of a Nilla wafer once a day starting at 8:00 with their assigned prenatal treatment group. This time period includes the start of brain sexual differentiation and when estrogen receptors begin to appear in the fetal brain (Wallen & Baum, 2002). Dams were weighed daily to calculate the volume of oil to pipette onto the wafer. All wafers were administered in separate cages without bedding that were used only by cage-pairs in the same prenatal treatment group. The dams readily ate the wafer with minimal handling by the experimenters. One dam refused the Nilla wafer following the first couple days of administration and was subsequently removed from the study. Experimenters involved with the administration of treatments were blinded to the treatment groups by using color codes for each group throughout the administration period.

The experimenters checked for births starting at GD 21 at 7:00 and 11:30. Pups were counted and sex-determined at 7:00 if the dam had finished labor (i.e., vaginal area cleaned, pups retrieved to the nest, dam nursing pups) and were considered postnatal day (PND) 0. Litters born between 7:00 and 11:30 were counted and sex-determined at 16:00. Pup sex was determined by relative anogenital distance. Litters with at least 6 male and 6 female pups were culled to 6 male and 6 female pups at PND 0 and kept for further testing. A total of 48 litters were used for postnatal testing (Corn Oil n = 12; 50 μg/kg BPA n = 11; 50 μg/kg Mixed BP n = 12; 150 μg/kg Mixed BP n = 13). Litters with an insufficient number of males or females (n = 43) were culled after birth and removed from the study. Initial litter size and sex ratio of pups at birth were recorded and analyzed from all litters born.

### 2.3 Postnatal Maternal Care Measures

#### 2.3.1 Video Recordings

Home cage maternal behavior was recorded using Raspberry Pi 3B+ minicomputers running Debian bullseye with the Raspberry Pi Desktop and equipped with Raspberry Pi Module 1 NoIR cameras. One Raspberry Pi was placed perpendicular to the short end and closest to the nest location for each cage. Raspberry Pis were programmed to record for one hour starting one hour after lights-off at 30 fps in greyscale at 1280 × 780 resolution from PND 1-10. Two infrared LED strip lights (940nm; LED Lights World) were attached to the bottom of the wire shelf above the cages and set to turn on and off for the recording automatically with a digital timer. Raspberry Pis were headless and accessed remotely in order to prevent disruption of home cage behavior by experimenter presence before or during the recordings. Following video recording, videos were automatically converted to mp4 format with MP4Box and uploaded to cloud storage. In the event that the dam moved the location of the nest to the opposite end of the cage, the camera side was also switched at the first opportunity. Full Raspberry Pi recording setup instructions and recording scripts are available at https://github.com/lapphe/raspberry_rat.

#### 2.3.2 Home Cage Behavior Analysis

Prior to analysis of maternal behavior, videos were screened to check that the camera was placed at the nest end of the cage and that the dam spent sufficient time on the nest. Of the 457 videos collected, 16 videos were not scorable for technical errors during recording (lighting issues, camera or cage moved during recording). In addition, the dam never entered the nest in 20 videos, the dam spent less than two minutes total on the nest in 95 videos, pups were at the wrong end of the cage in 74 videos, and 11 videos were of too low quality for automated scoring but could still be scorable manually. These 201 videos were manually scored according to the same ethogram used to generate the behavior classifiers in BORIS (Friard & Gamba, 2016). The remaining 240 videos were scored using the AMBER pipeline on a Dell Precision 7920 Tower with a Dual Intel Xeon Gold 5122 3.6GHz processor, 64GB RAM, Windows 10 operating system, and a NVIDIA Quadro P5000 video card (Lapp, Salazar, & Champagne, 2023). Probability thresholds used for behavior classifiers were 0.7 for nest attendance, 0.75 for nursing, and 0.45 for licking/grooming. Behavior annotations were filtered using a minimum bout length of 0.5 seconds for nest attendance, 2 seconds for nursing, and 0.25 seconds for licking/grooming. Total durations were calculated in SimBA for licking/grooming, nest attendance, active nursing, and passive nursing for each video (Nilsson et al., 2020).

All postnatal maternal care measures were normalized to seconds of observed behavior per day to account for missing observations. Passive nursing was observed infrequently and was therefore not used as an independent variable. Previous validation analyses have shown high agreement between manual scoring and AMBER scoring (Lapp, Salazar, & Champagne, 2023) and were not differentiated in downstream analyses for this study.

### 2.4 Pre-Weanling Measures

Pups were counted and weighed by sex at PND 0, 5, 10, 15, and 21. Eye-opening was assessed at PND 15 as a neurodevelopmental milestone. Pups were counted as having eyes open if they had at least one eye fully open (i.e., the lid membrane was entirely split open) when handled by the experimenter. Prevalence of eye-opening was then calculated as a percentage of all pups in the litter or split by sex. Brains were collected and flash frozen in hexanes on dry ice from up to one male pup and one female pup per litter at PND 10 and at PND 21 (weaning) so that four males and four females per litter were weaned for adult behavioral testing. Offspring were housed in same-sex sibling pairs at weaning and left undisturbed until young adulthood (PND 53+).

### 2.5 Adult Behavior Measures

#### 2.5.1 Attentional Set Shifting Task

A subset of offspring (n = 94) was tested for the attentional set shifting task using operant chambers as previously described (Brady & Floresco, 2015; Lauby et al., 2019). The attentional set shifting task assesses behavioral flexibility as the rat switches attention to a new dimension of stimuli while ignoring previously relevant stimuli (an attentional set; Brown & Tait, 2010). One male and one female sibling were tested from a total of 47 litters (Corn Oil n = 12; 50 μg/kg BPA n = 11; 50 μg/kg Mixed BP n = 12; 150 μg/kg Mixed BP n = 12). All operant sessions started at 12:00 during the dark phase and all offspring were acclimated to the testing room for at least 30 minutes prior to each session. To account for potential circadian rhythm effects, session times for offspring within each treatment group were counter-balanced between the six cohorts.

Sound-attenuating operant chambers (Habitest Modular System, Coulbourn Instruments) connected to a Dell desktop PC with Graphic State RT software (Coulbourn Instruments) installed were used for the attentional set shifting task. Each operant chamber was equipped with two retractable levers, two stimulus lights above the levers, a house light, and a food magazine that dispensed 45 mg banana sucrose pellets (#F0024, Bio-Serv).

All offspring (PND 53+) were food restricted to gradually reduce to 85-90% of their *ad libitum* feeding weight and handled two minutes per day for seven days before chamber acclimation. Food restriction was done to increase the motivational value of the sucrose pellet reward during training and testing in the operant chambers. Offspring received 20 sucrose pellets in their home cages the day before chamber acclimation. On the first session of operant testing, all offspring (PND 60+) were acclimated to the operant chamber with exposure to the house light for 30 minutes and 10 sucrose pellets dispensed (one pellet every 10 seconds) at the start of the acclimation session. Offspring completed an auto-shaping procedure for at least seven days to acquire the lever press response. Offspring with low rates of lever pressing (< 10 presses) by the seventh day of auto-shaping were manually shaped to lever press by an experimenter for one or two 30-minute sessions followed by three sessions of auto-shaping. Following auto-shaping training, offspring were required to press one of the levers extended within ten seconds at least 85 times out of 90 trials (omission task). After reaching criterion on the omission task, side preference was assessed as previously described by Brady and Floresco’s adaptation of the attentional set shifting task (Brady & Floresco, 2015).

For the first part of the attentional set shifting task, offspring were reinforced to press the lever on the illuminated side of the chamber (cued task) until they made 10 consecutive correct responses. The illuminated side was randomized between trials and was never illuminated more than two consecutive trials. Each session involved 150 trials and offspring were removed from the study if criterion was not met after three sessions (450 trials). The day after reaching criterion on the cued task, offspring were reinforced to press the lever opposite of their side preference regardless of which side was illuminated (response task) until they made 10 consecutive correct responses. Number of trials to reach criterion, adjusting for the number of trials with no responses, and percentage of trials where the rats responded correctly on the response task were used as measures of set-shifting ability and behavioral flexibility. Fewer trials to reach criterion and higher percentage of correct trials reflected better performance on the task. Estrous cycle stage was not assessed for the female offspring during this task; however, previous work has suggested that variability across the estrous cycle in rodents would not be substantial enough to change the interpretation of several other behavioral phenotypes (Beery, 2018; Levy et al., 2023).

One female offspring (50 μg/kg BPA) was removed from the task for not reaching criterion in the omission task after three sessions, one male offspring (Corn Oil) and two female offspring (50 μg/kg BPA and 150 μg/kg Mixed BP) were removed from the task for not reaching criterion in the cued task after 450 trials and one female offspring (150 μg/kg Mixed BP) was removed from analysis for completing the response task after 11 trials, leaving a total of 89 offspring reaching all criteria.

#### 2.5.2 Anxiety-Like Behavior

A separate subset of offspring were tested for anxiety-like behavior between PND 68-72, without prior handling. Male and female offspring were tested on different days approximately 3-6 hours after lights-off. Animals were habituated to the testing room for at least 15 minutes before each test. Arenas were cleaned with 70% ethanol solution and deionized water between tests. Estrous cycle stage was not assessed for the female offspring during these tasks. Pose estimation models and analysis scripts used to analyze the tests are available at https://github.com/lapphe/automated_anxiety.

##### 2.5.2.1 Elevated Plus Maze Test (EPM)

Up to two male and two female offspring from a total of 38 litters (Corn Oil n = 9; 50 μg/kg BPA n = 8; 50 μg/kg Mixed BP n = 10; 150 μg/kg Mixed BP n = 11) were tested in the EPM. The EPM was elevated 50 cm above the ground with 50 cm long arms enclosed by 40 cm walls and approximately 50-65 LUX in closed arms and 190-215 LUX in open arms. Offspring were placed in one of the closed arms of the maze and left to explore for 5 minutes. Offspring that fell off the maze before the end of the test (Corn Oil n = 2; 50 μg/kg BPA n = 4; 150 μg/kg Mixed BP n = 8) were excluded from analysis. EPM tests were video recorded from above and videos were analyzed using a DeepLabCut (DLC) pose estimation model generated in our lab to track 14 points on the rat and 12 points on the maze (Mathis et al., 2018). The EPM pose estimation model was trained on 695 labeled frames for 200,000 iterations and achieved a test error of 4.47 pixels on all points and 3.19 pixels on points with likelihood scores above 0.6. Points with likelihood scores of less than 0.5 were removed. Pose estimation data was then used with modified DLC analyzer code to extract time spent in the open arms and closed arms for each video (Sturman et al, 2020). Higher total time spent in the open arms and lower total time spent in the closed arms in the EPM reflected lower anxiety-like behavior. Offspring were considered in these zones using the location of the center back point (back_3).

##### 2.5.2.2 Open Field Test

Up to two male and two female offspring from a total of 48 litters (Corn Oil n = 12; 50 μg/kg BPA n = 11; 50 μg/kg Mixed BP n = 12; 150 μg/kg Mixed BP n = 13) were tested in the open field test. The open field was a 60 x 60 x 60 cm plexiglass testing arena with lighting of approximately 65-80 LUX. Offspring were placed in the middle of a testing arena and allowed to explore freely for 10 minutes. Behavior was recorded by a camera placed above the arena. Open field videos were analyzed using a DLC pose estimation model generated in our lab to track 14 points on the rat and 5 points on the arena (Mathis et al., 2018). The open field pose estimation model was trained on 585 labeled frames for 160,000 iterations and achieved a test error of 7.65 pixels on all points and 6.18 pixels on points with likelihood scores above 0.6. Pose estimation performance was also validated by visually inspecting labeled videos. Pose estimation data was then used with modified DLC analyzer code to extract time spent in the center and corners for each video (Sturman et al, 2020). Points with likelihood scores of less than 0.5 were removed. The center of the arena was set to the center 30 x 30 cm zone and the corners were defined as four 10 x 10 cm zones at the corners of the arena. Higher total time spent in the center and lower total time spent in the corners of the open field reflected lower anxiety-like behavior. Offspring were considered in these zones using the location of the center back point.

### 2.6 Brain Tissue Collection, RNA Extraction, and 3’ Tag-Sequencing

All brains were stored at -80° C until cryosectioning. PND 10 brains (n = 7 per sex per prenatal treatment group) were cryosectioned in 50 μm slices using a ThermoFisher Scientific CryoStar NX50 cryostat. The medial prefrontal cortex (+3.00 to +2.00 mm Bregma) and amygdala (-1.40 to -2.40 mm Bregma) were microdissected using an atlas for the developing rat brain (Paxinos et al., 1990) and a supplementary atlas for the PND 10 rat brain (Khazipov et al., 2015). RNA was extracted using the MagMAX mirVana Total RNA isolation kit with the Kingfisher Flex system according to manufacturer’s instructions (#A27828, ThermoFisher Scientific). RNA quantity was assessed with the Quant-iT RNA Assay kit (#Q33140, ThermoFisher Scientific) and RNA quality was assessed using the RNA 6000 Pico Assay kit (#5067-1513, Agilent Technologies). Two male medial prefrontal cortex samples (50 μg/kg Mixed BP and 150 μg/kg Mixed BP) were not collected during cryosectioning due to technical issues with the cryostat and one male medial prefrontal cortex sample (50 μg/kg Mixed BP) was lost during the DNase step for the RNA extractions.

All RNA samples were diluted to 25 ng/μl in 25 μl nuclease-free water before submission for library preparation and 3’ Tag-sequencing (Tag-seq; Lohman et al., 2016; Meyer et al., 2011) at the Genome Sequence and Analysis Facility at the University of Texas at Austin. Reads were sequenced on the NovaSeq S1 (100 bp single-end reads) with three to five million reads per sample.

### 2.7 Tag-seq Pipeline and Analysis

#### 2.7.1 Preprocessing

The tag-seq pipeline overview can be found in Figure 1B. Adaptor sequences, poly-A tails (> 8 basepairs long), and low-quality reads were trimmed using a custom script for tag-seq processing (https://github.com/z0on/tag-based_RNAseq) followed by cutadapt (version 1.18). Trimmed reads were mapped to the rat genome (*mRatBN7.2*) using STAR (version 2.7.3a) with the default settings (output files were unsorted BAM) and sorted using samtools (version 1.10). About 90% (+/- 1%) of reads were uniquely mapped to the rat genome for all samples. Gene counts were calculated for each sample using the bedtools (version 2.27.1) multicov function.

#### 2.7.2 Differential Gene Expression Analysis

To examine differential expression of individual genes, gene count data was normalized and analyzed for differential expression using the DESeq2 R package (version 1.34.0), separated by brain region and sex. Genes with more than 5 counts for more than 7 samples were analyzed. Prenatal Treatment and the postnatal maternal care measures (licking/grooming, nest attendance) from PND 1-5 were used as independent variables. Active nursing did not show significant main effects or interactions with prenatal treatment for the neurodevelopment or later-life behavior outcome measures and was therefore not included in any tag-seq analyses. Each postnatal maternal care measure was scaled and centered where the average is 0 and standard deviation is 1. Comparisons between prenatal treatment groups and interactions with prenatal treatment group and postnatal maternal care were always done relative to the Corn Oil group. Differentially expressed genes (DEGs) were identified with a false discovery rate (FDR) alpha ≤ 0.10 with Benjamani-Hochburg correction. All effects were reported as statistically significant if p ≤ 0.05 and marginally significant if p ≤ 0.10.

#### 2.7.3 Weighted Gene Co-expression Network Analysis (WGCNA)

The expression of genes can be highly correlated via interactions with overlapping transcription factors. To examine differential expression of co-expressing gene sets, gene networks were created with the WGCNA R package (version 1.72-1) using normalized and log-transformed gene count data, separated by brain region and sex. Genes with very low counts and variation were filtered prior to network construction using the goodSamplesGenes function in R with verbosity set to 3. WGCNA constructs matrices with Pearson correlations between every gene and creates clusters of highly correlated genes into eigengene modules (minimum size of 30 genes per module) using a soft power threshold of 8. A portion of modules had eigengene values that were characterized by one extreme outlier (> 0.90) and low variation following outlier removal (standard deviation < 0.10) and were therefore excluded from further analyses. The eigengene values for the remaining modules were analyzed using the lm function in R. Prenatal treatment and the postnatal maternal care measures (licking/grooming, nest attendance) from PND 1-5 were used as independent variables. Each postnatal maternal care measure was scaled and centered where the average is 0 and standard deviation is 1. Comparisons between prenatal treatment groups and interactions with prenatal treatment group and postnatal maternal care were always done relative to the Corn Oil group. For any eigengene module that showed statistically significant main effects or interactions, the gene sets were extracted and analyzed using Enrichr (Chen et al., 2013; Kuleshov et al., 2016; Xie et al., 2021) using the full gene list used for network construction as the background (https://maayanlab.cloud/Enrichr/). Enrichment from GO Biological Process 2023, Transcription Factor (TF) Protein-Protein Interactions, and ENCODE and ChEA Consensus TFs from ChIP-X were reported below. Transcription Factor Protein-Protein Interactions predict transcription factors that interact with the input gene set list (e.g., heterodimers, coactivators, corepressors, histone modifying enzymes). ENCODE and ChEA Consensus TFs from ChIP-X predict transcription factors that facilitate transcription of the input gene set. All effects were reported as statistically significant if p ≤ 0.05 and marginally significant if p ≤ 0.10.

### 2.8 Statistical Analysis

All statistical analyses were performed using R version 4.1.3 (R Core Team, 2023). Linear mixed models were performed using the lme4 R package and F-tests were performed using the ANOVA (Type III) function in the cars R package. Significant main effects of prenatal treatment were followed with t-tests between the Corn Oil group and each of the bisphenol treatment groups using the lmerTest R package. Batch of breeding was used as a random factor for all outcomes and litter ID was used as a random factor for the anxiety-like behavior measures to account for potential litter effects when testing multiple same-sex siblings on the same task. Distance traveled on the apparatus was used as a covariate for the anxiety-like behavior measures.

Two separate statistical models were used for the later-life outcome measures. To examine the main effects of prenatal bisphenol exposure alone on eye-opening, adult attentional set shifting and anxiety-like behavior, a linear mixed model was used with prenatal treatment and offspring sex as independent variables. To examine the interactive effects of prenatal bisphenol exposure and postnatal maternal care on eye-opening, adult attentional set shifting and anxiety-like behavior, a linear mixed model was used with prenatal treatment and the postnatal maternal care measures (licking/grooming, nest attendance, active nursing) as independent variables. Analyses were stratified by offspring sex with the exception of eye-opening prevalence at PND 15, where total percentage of pups in the litter with eyes open was also analyzed. Three litters (out of 48 total) were excluded from analyses (two from the Corn Oil group and one from the 150 μg/kg Mixed BP group) because they had more than two observations missing from the first five days of maternal behavior recordings. Analyses were based on postnatal maternal care from PND 1-5 since variations in maternal care in the first few postnatal days have been shown to have a substantial effect on neurodevelopment and behavior outcomes (Champagne et al., 2003; Hellstrom et al., 2012; Weaver et al., 2004).

Significant or marginal interactions of prenatal bisphenol treatment and postnatal maternal care were followed with a simple moderation analysis using PROCESS (Hayes, 2022) with the process function in R. A moderation tests for an interaction between an independent variable (*X;* Prenatal Treatment) and a moderator variable (*W;* Postnatal Maternal Care) on a dependent variable (*Y*). PROCESS is a flexible modelling module that can conduct moderation analyses and probe conditional effects of a focal moderator at the 16^th^, 50^th^, and 84^th^ percentile values (low, mid, and high postnatal maternal care respectively).

The statistical model used for a simple moderation (Model 1) was:

Conditional direct effects of *X* on *Y* = *intercept* + *b_1_X + b_2_W* + *b_3_X*W*

All effects were reported as statistically significant if p ≤ 0.05 and marginally significant if p ≤ 0.10.

## 3. Results

The full descriptions of the moderation, differential gene expression, and WGCNA analyses can be found in the supplementary results file.

### 3.1 Postnatal Maternal Care Moderates the Effects of Prenatal Bisphenol Exposure on Neurodevelopment

When examining the effects of prenatal treatment alone, there were no main effects of prenatal treatment or offspring sex or interactions between prenatal treatment and offspring sex on eye-opening prevalence at PND 15 (Figure 2A). When examining the interactive effects of prenatal treatment and postnatal maternal care, there was a marginal interaction between prenatal treatment and postnatal licking/grooming for total pups in the litter with eyes open (F_3,37_ = 2.5828, p = 0.06794; Figure 2B) and male pups in the litter with eyes open (F_3,37_ = 2.8123, p = 0.05264; Figure 2C) but not female pups in the litter with eyes open (F_3,37_ = 1.8762, p = 0.1505; Figure 2D). In addition, after accounting for postnatal licking/grooming and the interaction between prenatal treatment and postnatal licking/grooming, there was a main effect of prenatal treatment on total pups in the litter with eyes open (F_3,37_ = 2.9816, p = 0.04367; Figure 2B) and male pups in the litter with eyes open (F_3,37_ = 3.7856, p = 0.01828; Figure 2C). For total pups in the litter, there were significant differences between the Corn Oil group and the 50 μg/kg BPA (t = -2.048, p = 0.0477), 50 μg/kg Mixed BP (t = -2.623, p = 0.0126), and 150 μg/kg Mixed BP (t = -2.690, p = 0.0107) exposure groups, where prenatal bisphenol exposure was associated with reduced eye-opening prevalence at PND 15. For male pups in the litter, there were marginal differences between the Corn Oil group and the 50 μg/kg BPA (t = -1.868, p = 0.06972) exposure group and significant differences between the Corn Oil group and the 50 μg/kg Mixed BP (t = -3.272, p = 0.00232), and 150 μg/kg Mixed BP (t = -2.646, p = 0.01190) exposure groups, where prenatal bisphenol exposure was associated with reduced eye-opening prevalence for male pups at PND 15.

**Figure 2.**
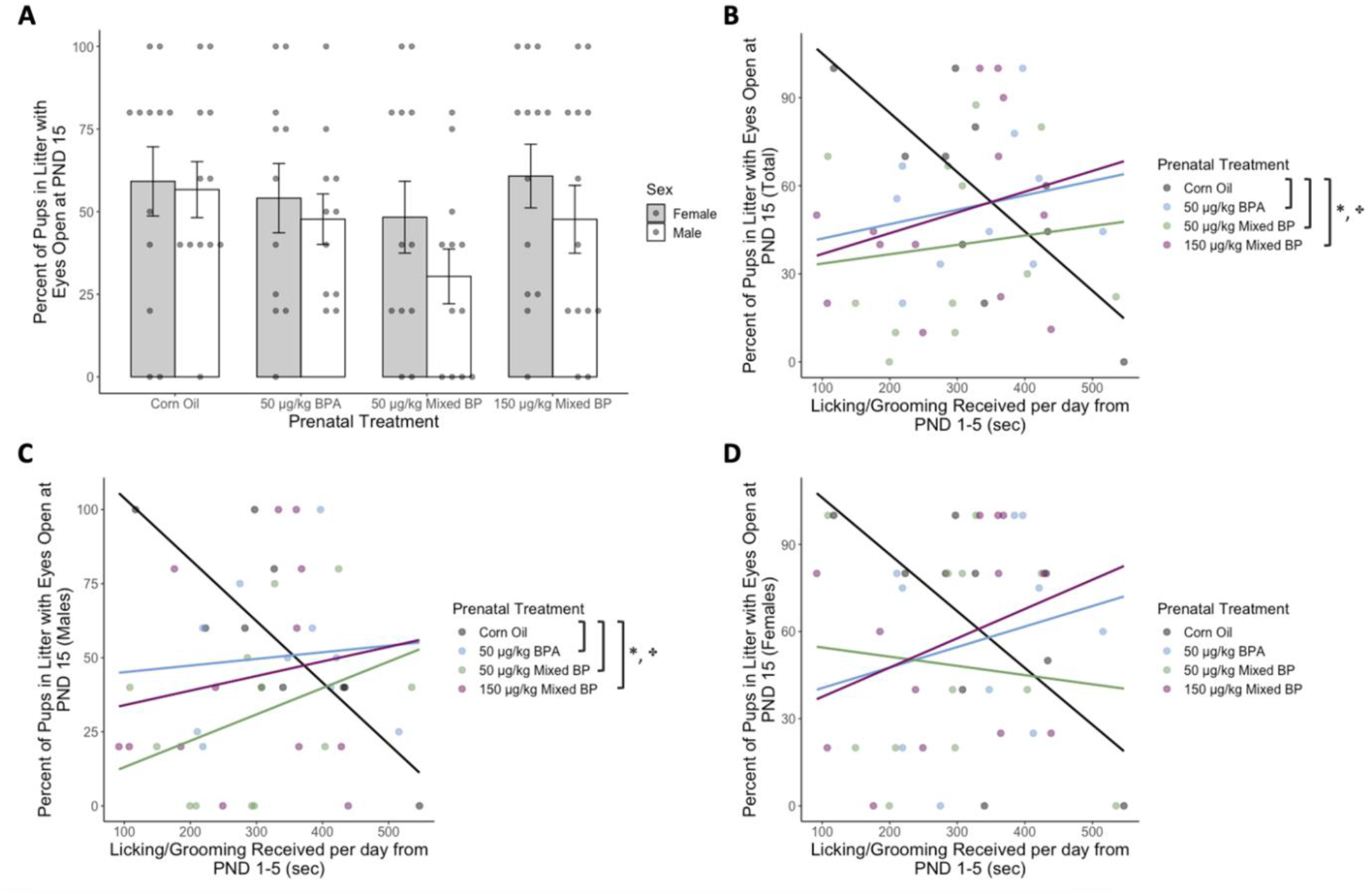
Prenatal bisphenol exposure was associated with reduced eye-opening prevalence at PND 15 after accounting for postnatal licking/grooming from PND 1-5, particularly in male pups. (A) No main effects of prenatal treatment were found for eye-opening prevalence at PND 15. Marginal interactions between prenatal treatment and postnatal licking/grooming were found for (B) total pups within a litter and (C) male pups only, but not (D) female pups only. After accounting for postnatal licking/grooming, all prenatal bisphenol exposure groups showed reduced eye-opening prevalence compared to the Corn Oil group for total pups within a litter and male pups only. Bar plots are displayed with mean +/- SEM with individual datapoints. Scatterplots are displayed with linear regression lines for each prenatal treatment group. * p < 0.05 main effect of prenatal treatment; ✤ p < 0.10 interaction between prenatal treatment and postnatal maternal care

Moderation analyses with PROCESS confirmed the marginal interactions between prenatal treatment and postnatal licking/grooming for total pups in the litter with eyes open (R^2^ change = 0.1636, F_3,37_ = 2.5828, p = 0.06790) and male pups in the litter with eyes open (R^2^ change = 0.1681, F_3,37_ = 2.8123, p = 0.0526). For offspring with low levels of licking/grooming, there was a reduction in prevalence in eye-opening with all prenatal BP exposure groups compared to the Corn Oil group (Table 1). There were no significant differences between prenatal treatment groups for offspring with high levels of licking/grooming. Similarly, for male offspring with low levels of licking/grooming, there was a reduction in prevalence in eye-opening with all prenatal BP exposure groups compared to the Corn Oil group (Table 1). There were no significant differences between prenatal treatment groups for male offspring with high levels of licking/grooming (Table 1).

**Table 1.**
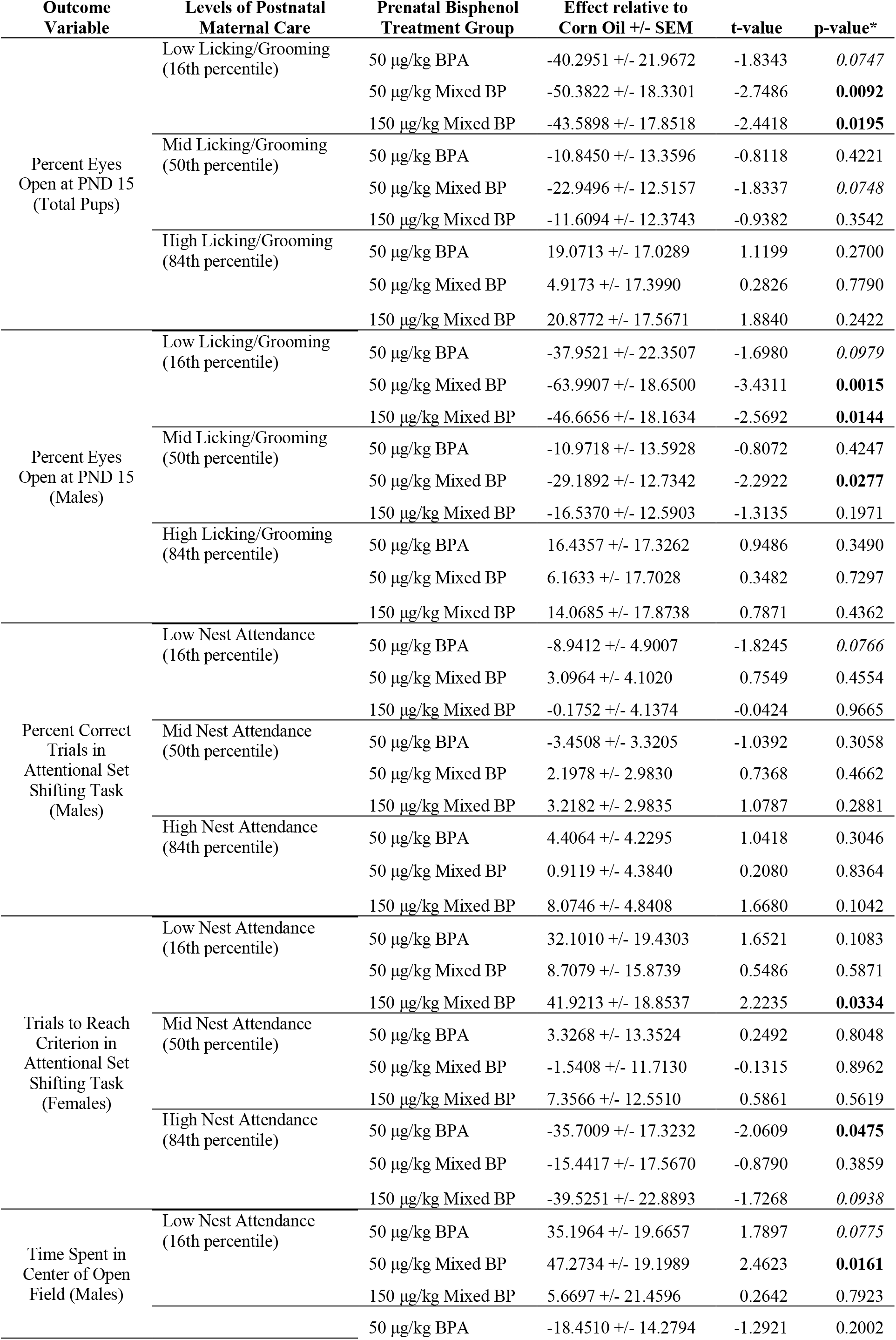
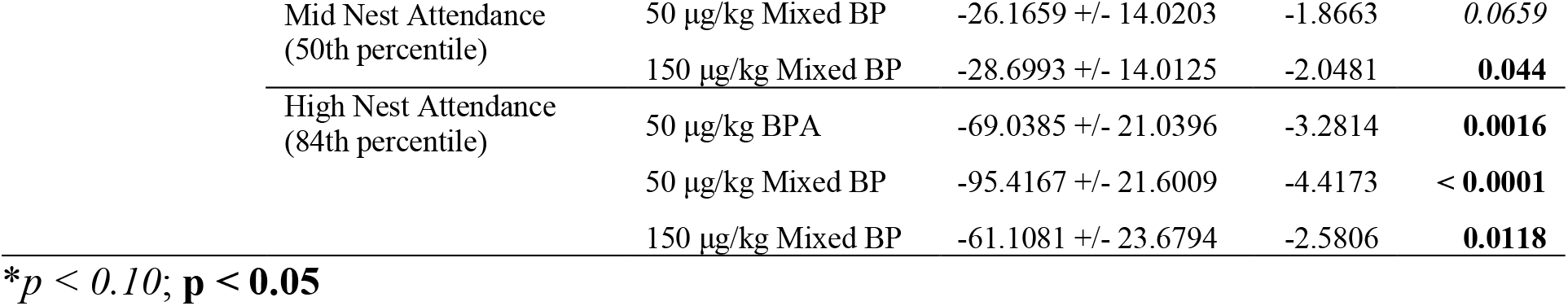
Summary of conditional post-hoc tests between the corn oil group and prenatal bisphenol treatment groups at the 16^th^, 50^th^, and 84^th^ values of postnatal maternal care.

| Outcome Variable | Levels of Postnatal Maternal Care | Prenatal Bisphenol Treatment Group | Effect relative to Corn Oil +/- SEM | t-value | p-value* |
| --- | --- | --- | --- | --- | --- |
| Percent Eyes Open at PND 15 (Total Pups) | Low Licking/Grooming (16th percentile) | 50 µg/kg BPA | -40.2951 +/- 21.9672 | -1.8343 | 0.0747 |
|  |  | 50 µg/kg Mixed BP | -50.3822 +/- 18.3301 | -2.7486 | **0.0092** |
|  |  | 150 µg/kg Mixed BP | -43.5898 +/- 17.8518 | -2.4418 | **0.0195** |
|  | Mid Licking/Grooming (50th percentile) | 50 µg/kg BPA | -10.8450 +/- 13.3596 | -0.8118 | 0.4221 |
|  |  | 50 µg/kg Mixed BP | -22.9496 +/- 12.5157 | -1.8337 | 0.0748 |
|  |  | 150 µg/kg Mixed BP | -11.6094 +/- 12.3743 | -0.9382 | 0.3542 |
|  | High Licking/Grooming (84th percentile) | 50 µg/kg BPA | 19.0713 +/- 17.0289 | 1.1199 | 0.2700 |
|  |  | 50 µg/kg Mixed BP | 4.9173 +/- 17.3990 | 0.2826 | 0.7790 |
|  |  | 150 µg/kg Mixed BP | 20.8772 +/- 17.5671 | 1.8840 | 0.2422 |
| Percent Eyes Open at PND 15 (Males) | Low Licking/Grooming (16th percentile) | 50 µg/kg BPA | -37.9521 +/- 22.3507 | -1.6980 | 0.0979 |
|  |  | 50 µg/kg Mixed BP | -63.9907 +/- 18.6500 | -3.4311 | **0.0015** |
|  |  | 150 µg/kg Mixed BP | -46.6656 +/- 18.1634 | -2.5692 | **0.0144** |
|  | Mid Licking/Grooming (50th percentile) | 50 µg/kg BPA | -10.9718 +/- 13.5928 | -0.8072 | 0.4247 |
|  |  | 50 µg/kg Mixed BP | -29.1892 +/- 12.7342 | -2.2922 | **0.0277** |
|  |  | 150 µg/kg Mixed BP | -16.5370 +/- 12.5903 | -1.3135 | 0.1971 |
|  | High Licking/Grooming (84th percentile) | 50 µg/kg BPA | 16.4357 +/- 17.3262 | 0.9486 | 0.3490 |
|  |  | 50 µg/kg Mixed BP | 6.1633 +/- 17.7028 | 0.3482 | 0.7297 |
|  |  | 150 µg/kg Mixed BP | 14.0685 +/- 17.8738 | 0.7871 | 0.4362 |
| Percent Correct Trials in Attentional Set Shifting Task (Males) | Low Nest Attendance (16th percentile) | 50 µg/kg BPA | -8.9412 +/- 4.9007 | -1.8245 | 0.0766 |
|  |  | 50 µg/kg Mixed BP | 3.0964 +/- 4.1020 | 0.7549 | 0.4554 |
|  |  | 150 µg/kg Mixed BP | -0.1752 +/- 4.1374 | -0.0424 | 0.9665 |
|  | Mid Nest Attendance (50th percentile) | 50 µg/kg BPA | -3.4508 +/- 3.3205 | -1.0392 | 0.3058 |
|  |  | 50 µg/kg Mixed BP | 2.1978 +/- 2.9830 | 0.7368 | 0.4662 |
|  |  | 150 µg/kg Mixed BP | 3.2182 +/- 2.9835 | 1.0787 | 0.2881 |
|  | High Nest Attendance (84th percentile) | 50 µg/kg BPA | 4.4064 +/- 4.2295 | 1.0418 | 0.3046 |
|  |  | 50 µg/kg Mixed BP | 0.9119 +/- 4.3840 | 0.2080 | 0.8364 |
|  |  | 150 µg/kg Mixed BP | 8.0746 +/- 4.8408 | 1.6680 | 0.1042 |
| Trials to Reach Criterion in Attentional Set Shifting Task (Females) | Low Nest Attendance (16th percentile) | 50 µg/kg BPA | 32.1010 +/- 19.4303 | 1.6521 | 0.1083 |
|  |  | 50 µg/kg Mixed BP | 8.7079 +/- 15.8739 | 0.5486 | 0.5871 |
|  |  | 150 µg/kg Mixed BP | 41.9213 +/- 18.8537 | 2.2235 | **0.0334** |
|  | Mid Nest Attendance (50th percentile) | 50 µg/kg BPA | 3.3268 +/- 13.3524 | 0.2492 | 0.8048 |
|  |  | 50 µg/kg Mixed BP | -1.5408 +/- 11.7130 | -0.1315 | 0.8962 |
|  |  | 150 µg/kg Mixed BP | 7.3566 +/- 12.5510 | 0.5861 | 0.5619 |
|  | High Nest Attendance (84th percentile) | 50 µg/kg BPA | -35.7009 +/- 17.3232 | -2.0609 | **0.0475** |
|  |  | 50 µg/kg Mixed BP | -15.4417 +/- 17.5670 | -0.8790 | 0.3859 |
|  |  | 150 µg/kg Mixed BP | -39.5251 +/- 22.8893 | -1.7268 | 0.0938 |
| Time Spent in Center of Open Field (Males) | Low Nest Attendance (16th percentile) | 50 µg/kg BPA | 35.1964 +/- 19.6657 | 1.7897 | 0.0775 |
|  |  | 50 µg/kg Mixed BP | 47.2734 +/- 19.1989 | 2.4623 | **0.0161** |
|  |  | 150 µg/kg Mixed BP | 5.6697 +/- 21.4596 | 0.2642 | 0.7923 |
|  |  | 50 µg/kg BPA | -18.4510 +/- 14.2794 | -1.2921 | 0.2002 |

|  |  |  |  |  |
| --- | --- | --- | --- | --- |
| Mid Nest Attendance<br>(50th percentile) | 50 µg/kg Mixed BP | -26.1659 +/- 14.0203 | -1.8663 | 0.0659 |
|  | 150 µg/kg Mixed BP | -28.6993 +/- 14.0125 | -2.0481 | **0.044** |
| High Nest Attendance<br>(84th percentile) | 50 µg/kg BPA | -69.0385 +/- 21.0396 | -3.2814 | **0.0016** |
|  | 50 µg/kg Mixed BP | -95.4167 +/- 21.6009 | -4.4173 | **< 0.0001** |
|  | 150 µg/kg Mixed BP | -61.1081 +/- 23.6794 | -2.5806 | **0.0118** |

*\*p* < 0.10; **p** < 0.05

### 3.3 Postnatal Maternal Care Moderates the Effects of Prenatal Bisphenol Exposure on Adult Behavior

#### 3.3.1 Attentional Set-Shifting Performance

When examining the effects of prenatal treatment alone, there were no main effects of prenatal treatment or offspring sex or interactions between prenatal treatment and offspring sex on trials to reach criterion (Figure 3A) or percent correct trials (Figure 3B) on the attentional set shifting task. When examining the interactive effects of prenatal treatment and postnatal maternal care, there was a marginal interaction between prenatal treatment and postnatal nest attendance for percent correct trials in male offspring (F_3,35_ = 2.8063, p = 0.05386; Figure 3C) and a significant interaction between prenatal treatment and postnatal nest attendance for trials to reach criterion in female offspring (F_3,32_ = 3.3530, p = 0.03095; Figure 3D). In addition, after accounting for postnatal nest attendance and the interaction between prenatal treatment and postnatal nest attendance, there was a main effect of prenatal treatment on percent correct trials in male offspring (F_3,35_ = 2.9017, p = 0.04852) and trials to reach criterion in female offspring (F_3,32_ = 3.3530, p = 0.03095). For the percent correct trials in male offspring, there was a significant difference between the Corn Oil group and the 50 μg/kg BPA (t = -2.073, p = 0.0456) exposure group, where prenatal BPA exposure was associated with a reduced percentage of correct trials. For the trials to reach criterion in female offspring, there were significant differences between the Corn Oil group and the 50 μg/kg BPA (t = -2.442, p = 0.0203) and 150 μg/kg Mixed BP (t = 2.518, p = 0.0170) exposure groups, where prenatal bisphenol exposure was associated with an increased number of trials to reach criterion.

**Figure 3.**
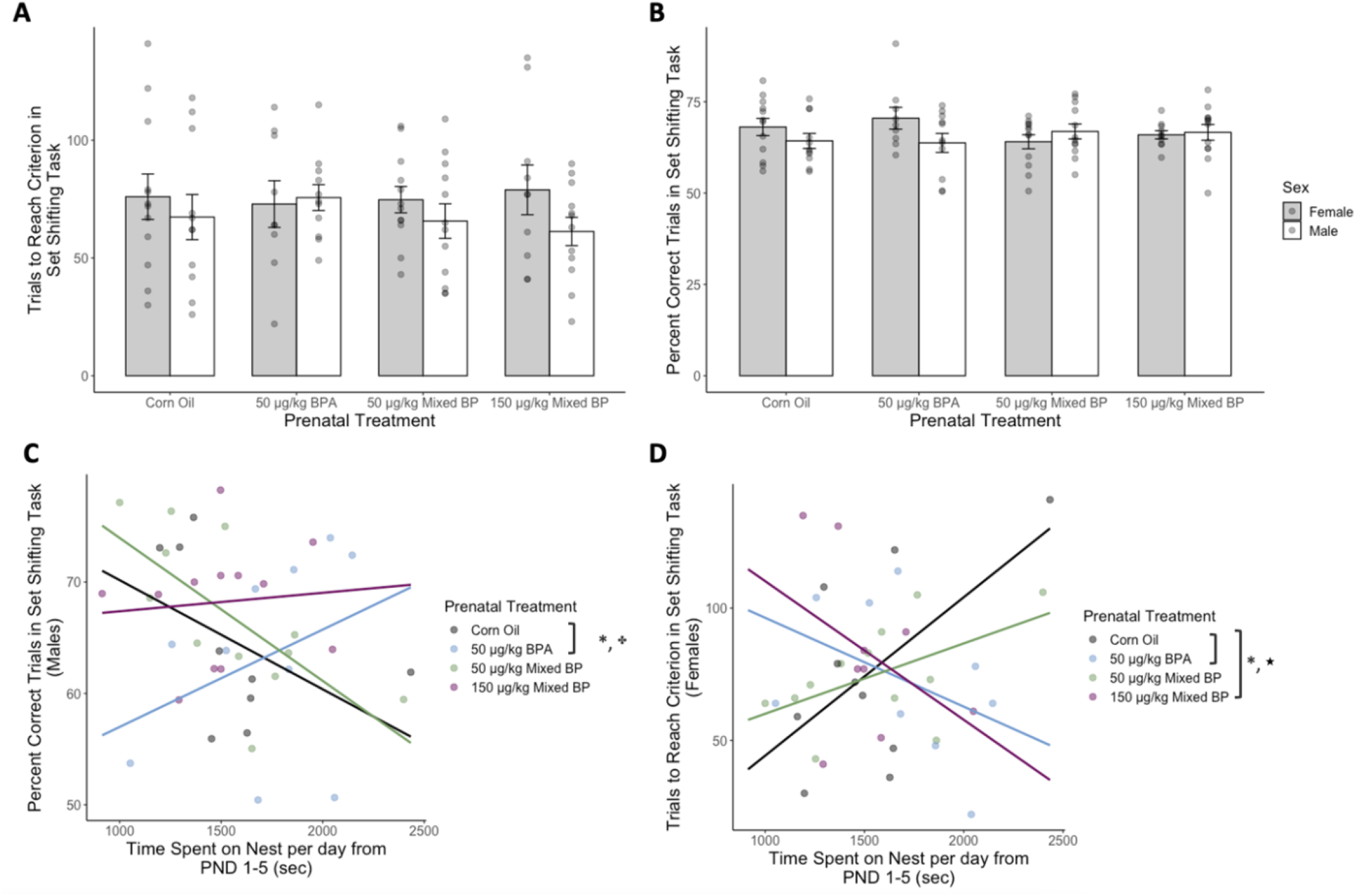
Prenatal bisphenol exposure was associated with reduced attentional set shifting performance after accounting for postnatal nest attendance from PND 1-5 for male and female offspring. No main effects of prenatal treatment were found for (A) trials to reach criterion or (B) percent correct trials in the attentional set shifting response task. A marginal interaction between prenatal treatment and postnatal nest attendance was found for (C) percent correct trials for male offspring. A significant interaction between prenatal treatment and postnatal nest attendance was found for (D) trials to reach criterion for female offspring. After accounting for postnatal nest attendance, the 50 μg/kg BPA (male and female offspring) and 150 μg/kg Mixed BP (female offspring only) exposure groups showed reduced attentional set shifting performance compared to the Corn Oil group. Bar plots are displayed with mean +/- SEM with individual datapoints. Scatterplots are displayed with linear regression lines for each prenatal treatment group. * p < 0.05 main effect of prenatal treatment; ★ p < 0.05 interaction between prenatal treatment and postnatal maternal care; ✤ p < 0.10 interaction between prenatal treatment and postnatal maternal care

Moderation analyses with PROCESS confirmed the marginal interaction between prenatal treatment and postnatal nest attendance for percent correct trials in male offspring (R^2^ change = 0.1725, F_3,35_ = 2.8063, p = 0.0539) and the significant interaction between prenatal treatment and postnatal nest attendance for trials to reach criterion in female offspring (R^2^ change = 0.2308, F_3,32_ = 3.3530, p = 0.0310). For male offspring with low levels of nest attendance, there was a reduction in percent correct trials with the 50 μg/kg BPA exposure group compared to the Corn Oil group (Table 1). There were no significant differences between prenatal treatment groups for male offspring with high levels of nest attendance (Table 1). For female offspring with low levels of nest attendance, there was an increase in the number of trials to reach criterion with the 150 μg/kg Mixed BP exposure group compared to the Corn Oil group. For female offspring with high levels of nest attendance, there was a reduction in the number of trials to reach criterion with the 50 μg/kg BPA and 150 μg/kg Mixed BP exposure group compared to the Corn Oil (Table 1).

#### 3.3.2 Anxiety-Like Behavior

When examining the effects of prenatal treatment alone, there were no main effects of prenatal treatment or offspring sex on time spent in the open arms (Figure 4A) or closed arms (Figure 4B) in the elevated plus maze (EPM). There was a significant interaction between prenatal treatment and offspring sex on time spent in the closed arms in the EPM (F_3,85.262_ = 2.9418, p = 0.0376292; Figure 4B). There were no known main effects or interactions with postnatal maternal care.

**Figure 4.**
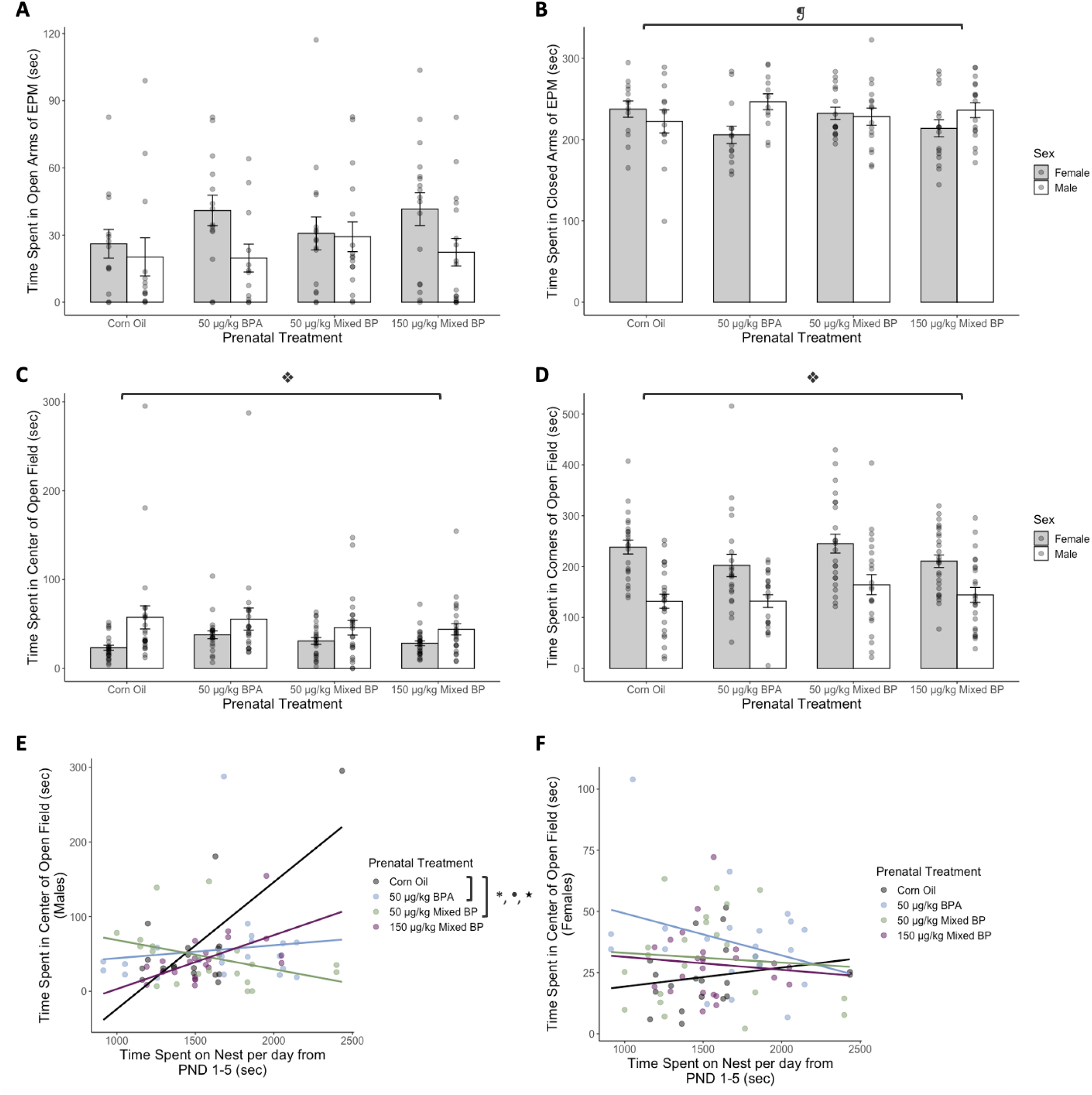
Prenatal bisphenol exposure was associated with reduced anxiety-like behavior in the open field after accounting for postnatal nest attendance from PND 1-5 for male offspring. No main effects of prenatal treatment were found for (A) time spent in open arms or (B) time spent in the closed arms of the elevated plus maze (EPM), although there was a significant interaction of prenatal treatment and offspring sex on time spent in the closed arms of the EPM. Significant main effects of offspring sex were found for (C) time spent in the center and (D) time spent in the corners of the open field, where female offspring were more anxious than male offspring. (E) A significant main effect of postnatal nest attendance was found for time spent in the center in the open field for male offspring. A significant interaction between prenatal treatment and postnatal nest attendance was found for time spent in the center in the open field for (E) male offspring but not (F) female offspring. After accounting for postnatal nest attendance, the 50 μg/kg BPA and 50 μg/kg Mixed BP exposure groups showed reduced anxiety-like behavior in the open field compared to the Corn Oil group for male offspring only. Bar plots are displayed with mean +/- SEM with individual datapoints. Scatterplots are displayed with linear regression lines for each prenatal treatment group. * p < 0.05 main effect of prenatal treatment; ❡ p < 0.05 interaction between prenatal treatment and offspring sex; ❖ p < 0.05 main effect of offspring sex; ● p < 0.05 main effect of postnatal maternal care; ★ p < 0.05 interaction between prenatal treatment and postnatal maternal care

When examining the effects of prenatal treatment alone, there were no main effects of prenatal treatment on time spent in the center or corners of the open field. There was a main effect of offspring sex on time spent in the center (F_1,170.42_ = 15.2308, p < 0.001; Figure 4C) and corners (F_1,132.497_ = 67.7646, p < 0.001; Figure 4D) of the open field and no interactions between prenatal treatment and offspring sex. Female offspring spent less time in the center (t = -3.261, p = 0.00134) and more time in the corners (t = 5.370, p < 0.001) in the open field than male offspring. When examining the interactive effects of prenatal treatment and postnatal maternal care, there was a significant interaction between prenatal treatment and postnatal nest attendance for time spent in the center of the open field for male offspring (F_3,75_ = 7.9383, p < 0.001; Figure 4E) but not female offspring (F_3,28.354_ = 0.9691, p = 0.420989; Figure 4F). There was also a significant main effect of postnatal nest attendance for time spent in the center of the open field for male offspring (F_1,75_ = 12.4024, p < 0.001; Figure 4E), where higher postnatal nest attendance was associated with increased time spent in the center. Finally, after accounting for postnatal nest attendance and the interaction between prenatal treatment and postnatal nest attendance, there was a main effect of prenatal treatment on time spent in the center of the open field for male offspring (F_3,75_ = 6.9481, p < 0.001; Figure 4E). There were significant differences between the Corn Oil group and the 50 μg/kg BPA (t = 3.264, p = 0.001657) and 50 μg/kg Mixed BP (t = 4.380, p < 0.001) exposure groups, where prenatal bisphenol exposure was associated with increased time spent in the center in the open field for male offspring.

Moderation analyses with PROCESS confirmed the significant interaction between prenatal treatment and postnatal nest attendance for time spent in the center of the open field for male offspring (R^2^ change = 0.2273, F_3,76_ = 8.1170, p = < 0.001). For male offspring with low levels of nest attendance, there was an increase in time spent in the center of the open field with the 50 μg/kg BPA and 50 μg/kg Mixed BP exposure groups compared to the Corn Oil group (Table 1). For male offspring with high levels of nest attendance, there was a significant reduction in time spent in the center of the open field with all the prenatal BP exposure groups compared to the Corn Oil group (Table 1).

### 3.4 Interactive Effects of Prenatal Bisphenol Exposure and Postnatal Maternal Care on PND 10 Brain Gene Expression

The summary of the differential gene expression analysis for male and female medial prefrontal cortex and amygdala can be found in Table 2. The results from the full statistical model (including prenatal treatment group and both postnatal maternal care measures) are reported. There were no DEGs identified between prenatal treatment groups without accounting for postnatal maternal care.

**Table 2.** Summary of differentially expressed genes compared to the corn oil group.

| Medial Prefrontal Cortex |  |  |  |  |  |  |
| --- | --- | --- | --- | --- | --- | --- |
|  | FDR < 0.1 |  |  | FDR < 0.05 |  |  |
|  | Direction of Effect |  |  | Direction of Effect |  |  |
| *Male* | Positive | Negative | Total | Positive | Negative | Total |
| 50 µg/kg BPA | 0 | 0 | 0 | 0 | 0 | 0 |
| 50 µg/kg Mixed BP | 0 | 0 | 0 | 0 | 0 | 0 |
| 150 µg/kg Mixed BP | 0 | 2 | 2 | 0 | 2 | 2 |
| Nest Attendance | 0 | 25 | 25 | 0 | 6 | 6 |
| Licking/Grooming | 0 | 0 | 0 | 0 | 0 | 0 |
| 50 µg/kg BPA x Nest Attendance | 0 | 0 | 0 | 0 | 0 | 0 |
| 50 µg/kg Mixed BP x Nest Attendance | 24 | 0 | 24 | 8 | 0 | 8 |
| 150 µg/kg Mixed BP x Nest Attendance | 22 | 3 | 25 | 6 | 0 | 6 |
| 50 µg/kg BPA x Licking/Grooming | 0 | 0 | 0 | 0 | 0 | 0 |
| 50 µg/kg Mixed BP x Licking/Grooming | 0 | 0 | 0 | 0 | 0 | 0 |
| 150 µg/kg Mixed BP x Licking/Grooming | 0 | 0 | 0 | 0 | 0 | 0 |
| *Female* |  |  |  |  |  |  |
| 50 µg/kg BPA | 0 | 0 | 0 | 0 | 0 | 0 |
| 50 µg/kg Mixed BP | 0 | 0 | 0 | 0 | 0 | 0 |
| 150 µg/kg Mixed BP | 0 | 1 | 1 | 0 | 0 | 1 |
| Nest Attendance | 0 | 1 | 1 | 0 | 0 | 1 |
| Licking/Grooming | 0 | 0 | 0 | 0 | 0 | 0 |
| 50 µg/kg BPA x Nest Attendance | 0 | 0 | 0 | 0 | 0 | 0 |
| 50 µg/kg Mixed BP x Nest Attendance | 0 | 1 | 1 | 0 | 1 | 1 |
| 150 µg/kg Mixed BP x Nest Attendance | 0 | 1 | 1 | 0 | 1 | 1 |
| 50 µg/kg BPA x Licking/Grooming | 0 | 0 | 0 | 0 | 0 | 0 |
| 50 µg/kg Mixed BP x Licking/Grooming | 0 | 0 | 0 | 0 | 0 | 0 |
| 150 µg/kg Mixed BP x Licking/Grooming | 0 | 0 | 0 | 0 | 0 | 0 |
| Amygdala |  |  |  |  |  |  |
| *Male* | Positive | Negative | Total | Positive | Negative | Total |
| 50 µg/kg BPA | 0 | 0 | 0 | 0 | 0 | 0 |
| 50 µg/kg Mixed BP | 2 | 285 | 287 | 0 | 0 | 0 |
| 150 µg/kg Mixed BP | 1 | 242 | 243 | 0 | 127 | 128 |
| Nest Attendance | 0 | 0 | 0 | 0 | 0 | 0 |
| Licking/Grooming | 0 | 0 | 0 | 0 | 0 | 0 |
| 50 µg/kg BPA x Nest Attendance | 0 | 0 | 0 | 0 | 0 | 0 |
| 50 µg/kg Mixed BP x Nest Attendance | 17 | 223 | 240 | 1 | 52 | 53 |
| 150 µg/kg Mixed BP x Nest Attendance | 16 | 500 | 516 | 10 | 340 | 350 |
| 50 µg/kg BPA x Licking/Grooming | 0 | 0 | 0 | 0 | 0 | 0 |
| 50 µg/kg Mixed BP x Licking/Grooming | 0 | 0 | 0 | 0 | 0 | 0 |
| 150 µg/kg Mixed BP x Licking/Grooming | 0 | 0 | 0 | 0 | 0 | 0 |

| *Female* |  |  |  |  |  |  |
| --- | --- | --- | --- | --- | --- | --- |
| 50 µg/kg BPA | 0 | 0 | 0 | 0 | 0 | 0 |
| 50 µg/kg Mixed BP | 0 | 0 | 0 | 0 | 0 | 0 |
| 150 µg/kg Mixed BP | 0 | 1 | 1 | 0 | 1 | 1 |
| Nest Attendance | 0 | 0 | 0 | 0 | 0 | 0 |
| Licking/Grooming | 0 | 0 | 0 | 0 | 0 | 0 |
| 50 µg/kg BPA x Nest Attendance | 0 | 0 | 0 | 0 | 0 | 0 |
| 50 µg/kg Mixed BP x Nest Attendance | 0 | 1 | 1 | 0 | 1 | 1 |
| 150 µg/kg Mixed BP x Nest Attendance | 0 | 0 | 0 | 0 | 0 | 0 |
| 50 µg/kg BPA x Licking/Grooming | 0 | 0 | 0 | 0 | 0 | 0 |
| 50 µg/kg Mixed BP x Licking/Grooming | 0 | 1 | 1 | 0 | 1 | 1 |
| 150 µg/kg Mixed BP x Licking/Grooming | 0 | 0 | 0 | 0 | 0 | 0 |

#### 3.4.1 Medial Prefrontal Cortex

##### 3.4.1.1 Male Offspring

There were significant interactions between the 50 μg/kg Mixed BP and 150 μg/kg Mixed BP exposure groups and postnatal nest attendance (Table 2). There were 24 DEGs with a positive R^2^ change for the 50 μg/kg Mixed BP exposure group and nest attendance compared to the Corn Oil group (Supplementary Table 2). There were 22 DEGs with a positive R^2^ change and 3 DEGS with a negative R^2^ change for the 150 μg/kg Mixed BP exposure group and nest attendance compared to the Corn Oil group (Supplementary Table 3). Some common DEGs with a positive R^2^ change included Nrgn (Supplementary Figure 2A), Tuba4a (Supplementary Figure 2B), Nefl (Supplementary Figure 2C), and Mt3 (Supplementary Figure 2D) (all FDR < 0.05).

A total of 10 eigengene modules were examined for differences between prenatal treatments and postnatal maternal care and their interactions. There were significant main effects or interactions found with five eigengene modules (Black, Brown, Lightcyan, Tan, and Yellow).

For the black module, there were significant interactions between the 50 μg/kg BPA exposure group and nest attendance (t = 2.170, p = 0.04443), 50 μg/kg Mixed BP exposure group and nest attendance (t = 2.662, p = 0.01641), and the 150 μg/kg Mixed BP exposure group and nest attendance (t = 2.242, p = 0.03861) compared to the Corn Oil group (Figure 5A). The top GO terms for the genes in the black module were Macromolecule Biosynthetic Process, Translation, Peptide Biosynthetic Process, Gene Expression, Regulation of Apoptotic Process, and Protein Transport. The top enriched transcription factors involved in protein-protein interactions with genes in the black module were MYC, ESR1, TP53, ILF3, and ATF2 (Figure 5A). The top enriched transcription factors involved in facilitating transcription of genes in the black module were GABPA, ELF1, TAF1, NFYB, and YY1 (Figure 5A).

**Figure 5.**
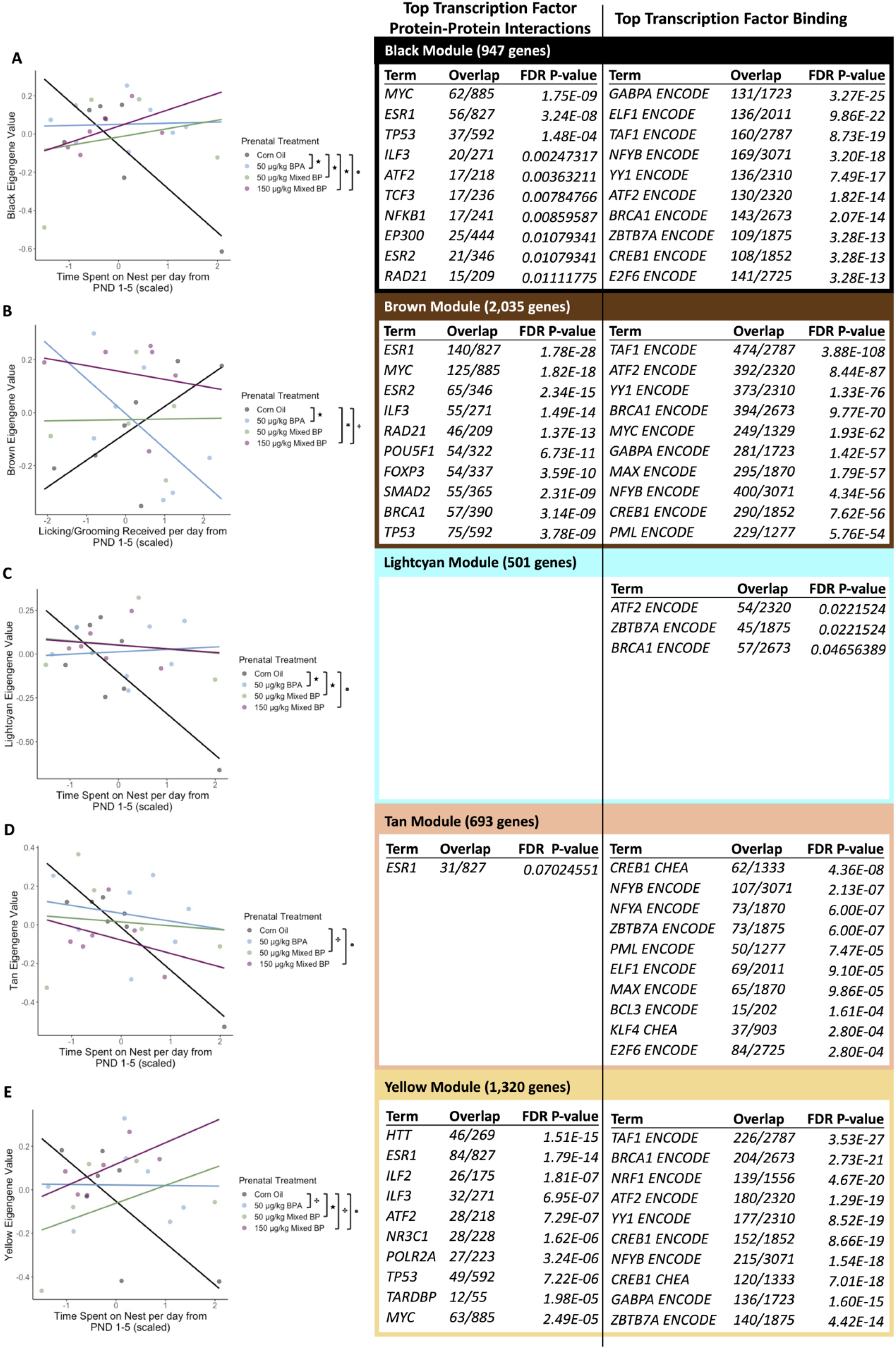
The interactive effects of prenatal bisphenol exposure and postnatal maternal care on transcriptomic profiles in the male medial prefrontal cortex. There were significant main effects or interactions with prenatal treatment and postnatal maternal care for the (A) black module, (B) brown module, (C) lightcyan module, (D) tan module, and (E) yellow module. Scatterplots are displayed with linear regression lines for each prenatal treatment group. ● p < 0.05 main effect of postnatal maternal care; + p < 0.10 main effect of postnatal maternal care; ★ p < 0.05 interaction between prenatal treatment and postnatal maternal care; ✤ p < 0.10 interaction between prenatal treatment and postnatal maternal care

For the brown module, there was a significant interaction between the 50 μg/kg BPA exposure group and licking/grooming (t = -2.704, p = 0.0150) compared to the Corn Oil group (Figure 5B). The top GO terms for the genes in the brown module were Translation, Gene Expression, Cytoplasmic Translation, and Peptide Biosynthetic Process. The top enriched transcription factors involved in protein-protein interactions with genes in the brown module were ESR1, MYC, ESR2, ILF3, and RAD21 (Figure 5B). The top enriched transcription factors involved in facilitating transcription of genes in the brown module were TAF1, ATF2, YY1, BRCA1, and MYC (Figure 5B).

For the lightcyan module, there were significant interactions between the 50 μg/kg BPA exposure group and nest attendance (t = 2.418, p = 0.02714) and the 50 μg/kg Mixed BP exposure group and nest attendance (t = 2.174, p = 0.04415) compared to the Corn Oil group (Figure 5C). There were no significantly enriched GO terms or transcription factor protein-protein interactions for the genes in the lightcyan module. The top enriched transcription factors involved in facilitating transcription of genes in the lightcyan module were ATF2, ZBTB7A, and BRCA1 (Figure 5C).

For the tan module, there was a marginal interaction between the 50 μg/kg Mixed BP exposure group and nest attendance (t = 1.917, p = 0.0722) compared to the Corn Oil group (Figure 5D). There were no significantly enriched GO terms and the top transcription factor protein-protein interaction, ESR1, was marginally significant (FDR = 0.0702) for the genes in the tan module (Figure 5D). The top enriched transcription factors involved in facilitating transcription of genes in the tan module were CREB1, NFYB, NFYA, ZBTB7A, and PML (Figure 5D).

For the yellow module, there was a significant interaction between the 50 μg/kg Mixed BP exposure group and nest attendance (t = 2.667, p = 0.0162) and marginal interactions between the 50 μg/kg BPA exposure group and nest attendance (t = 1.750, p = 0.0981) and the 150 μg/kg Mixed BP exposure group and nest attendance (t = 2.069, p = 0.0541) compared to the Corn Oil group (Figure 5E). The top GO terms for the genes in the yellow module were Proton Motive Force-Driven ATP Synthesis, Proton Motive Force-Driven Mitochondrial ATP Synthesis, Oxidative Phosphorylation, Signal Release from Synapse, and Neurotransmitter Secretion. The top enriched transcription factors involved in protein-protein interactions with genes in the yellow module were HTT, ESR1, ILF2, ILF3, and ATF2 (Figure 5E). The top enriched transcription factors involved in facilitating transcription of genes in the yellow module were TAF1, BRCA1, NRF1, ATF2, and YY1 (Figure 5E).

##### 3.4.1.2 Female Offspring

There were significant interactions between the 50 μg/kg BPA and 150 μg/kg Mixed BP exposure groups and postnatal nest attendance (Table 2). There was one DEG with a negative R^2^ change for the 50 μg/kg BPA exposure group and nest attendance compared to the Corn Oil group, which included Glra3 (Supplementary Figure 4; FDR < 0.05). There was one DEG with a negative R^2^ change for the 150 μg/kg BPA exposure group and nest attendance compared to the Corn Oil group, which included Dcx (FDR < 0.05). Glra3 was listed as the second top gene by unadjusted p-value but did not pass FDR p-value correction (FDR = 0.515).

A total of nine eigengene modules were examined for differences between prenatal treatments and postnatal maternal care and their interactions. There were significant main effects or interactions found with two eigengene modules (Brown and Turquoise).

For the brown module, there were significant interactions between the 50 μg/kg BPA exposure group and licking/grooming (t = -2.141, p = 0.04478) and the 50 μg/kg Mixed BP exposure group and licking/grooming (t = -3.431, p = 0.00265) compared to the Corn Oil group (Figure 6A). The top GO terms for the genes in the brown module were Cytoplasmic Translation, Macromolecule Biosynthetic Process, Gene Expression, and Translation. The top enriched transcription factors involved in protein-protein interactions with genes in the brown module were ESR1, ILF3, RAD21, POU5F1, and MYC (Figure 6A). The top enriched transcription factors involved in facilitating transcription of genes in the brown module were TAF1, YY1, ATF2, BRCA1, and MYC (Figure 6A).

**Figure 6.**
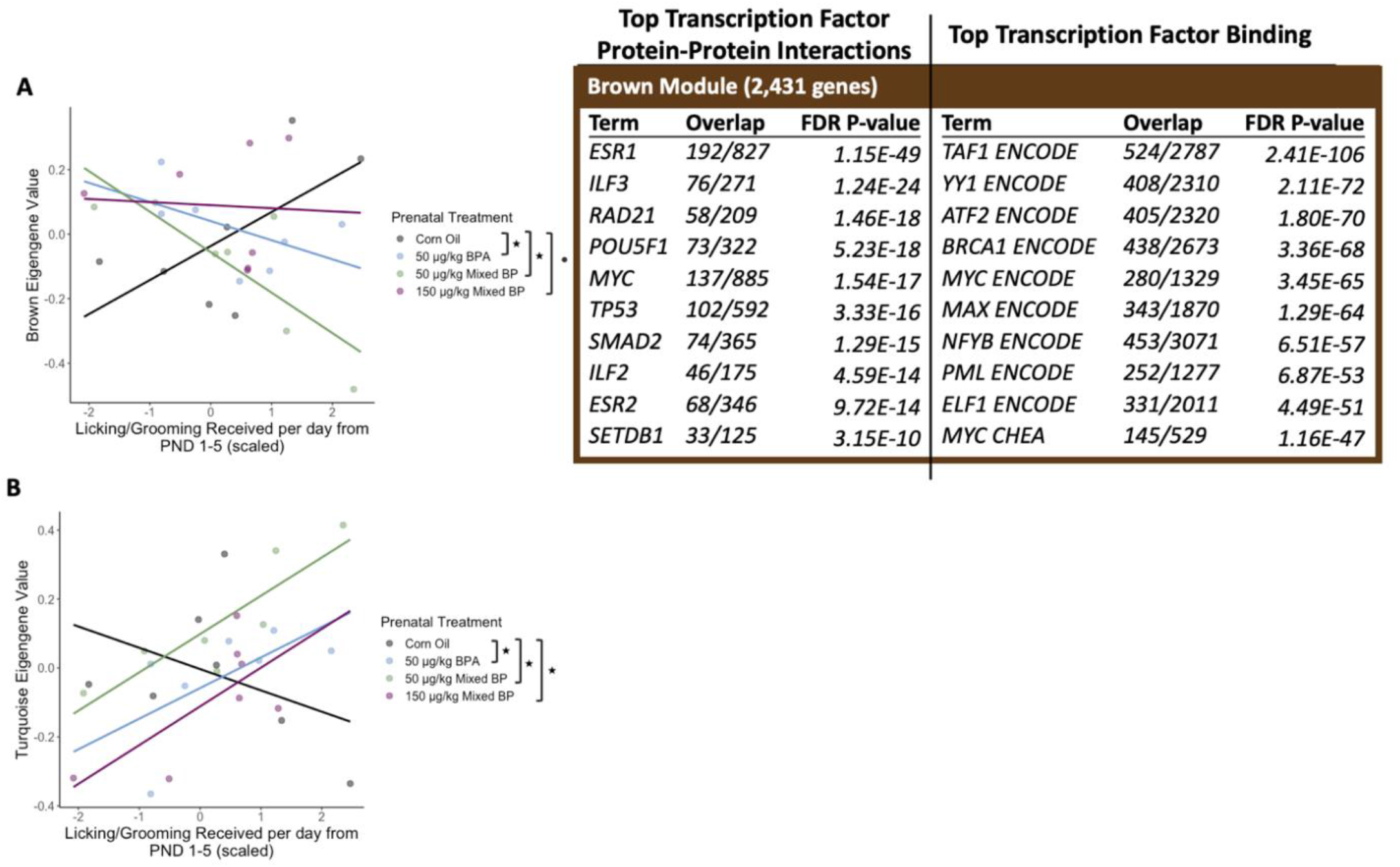
The interactive effects of prenatal bisphenol exposure and postnatal maternal care on transcriptomic profiles in the female medial prefrontal cortex. There were significant main effects or interactions with prenatal treatment and postnatal maternal care for the (A) brown module and (B) turquoise module. Scatterplots are displayed with linear regression lines for each prenatal treatment group. ● p < 0.05 main effect of postnatal maternal care; ★ p < 0.05 interaction between prenatal treatment and postnatal maternal care

For the turquoise module, there were significant interactions between the 50 μg/kg BPA exposure group and licking/grooming (t = 2.102, p = 0.0484), the 50 μg/kg Mixed BP exposure group and licking/grooming (t = 2.758, p = 0.0121), and the 150 μg/kg Mixed BP exposure group and licking/grooming (t = 2.452, p = 0.0235) compared to the Corn Oil group (Figure 6B). There were no significantly enriched GO terms, transcription factor protein-protein interactions, or transcription factors that facilitate transcription of the genes in the turquoise module.

#### 3.4.2 Amygdala

##### 3.4.2.1 Male Offspring

There were significant interactions between the 50 μg/kg Mixed BP and 150 μg/kg Mixed BP exposure groups and postnatal maternal care (Table 2). There were 17 DEGs with a positive R^2^ change and 223 DEGs with a negative R^2^ change for the 50 μg/kg Mixed BP exposure group and nest attendance compared to the Corn Oil group (Supplementary Table 6). There were 16 DEGs with a positive R^2^ change and 500 DEGs with a negative R^2^ change for the 150 μg/kg Mixed BP exposure group and nest attendance compared to the Corn Oil group (Supplementary Table 7). Some common DEGs with a negative R^2^ change included Tafa2 (Supplementary Figure 6A), Grip1 (Supplementary Figure 6B), Robo1 (Supplementary Figure 6C), and Epha3 (Supplementary Figure 6D; all FDR < 0.05). In addition, there was one DEG with a positive R^2^ change and one DEG with a negative R^2^ change for the 50 μg/kg Mixed BP exposure group and licking/grooming compared to the Corn Oil group, which included Cdh13 and Col1a2 respectively (Table 2; all FDR < 0.10).

A total of 14 eigengene modules were examined for differences between prenatal treatments and postnatal maternal care and their interactions. There were significant main effects or interactions found with two eigengene modules (Blue and Green).

For the blue module, there was a significant interaction between the 150 μg/kg Mixed BP exposure group and nest attendance (t = -3.226, p = 0.00423) and a marginal interaction between the 50 μg/kg Mixed BP exposure group and nest attendance (t = -1.733, p = 0.09849) compared to the Corn Oil group (Figure 7A). The top GO terms for the genes in the blue module were Modulation of Chemical Synaptic Transmission, Chemical Synaptic Transmission, Nervous System Development, Regulation of Transcription by RNA Polymerase II, Anterograde Trans-Synaptic Signaling and Axonogenesis. The top enriched transcription factors involved in protein-protein interactions with genes in the blue module were CTNNB1, MYC, SMAD2, FOXP3, and ESR1 (Figure 7A). The top enriched transcription factors involved in facilitating transcription of genes in the blue module were AR, UBTF, TAF1, SUZ12, and SMAD4 (Figure 7A).

**Figure 7.**
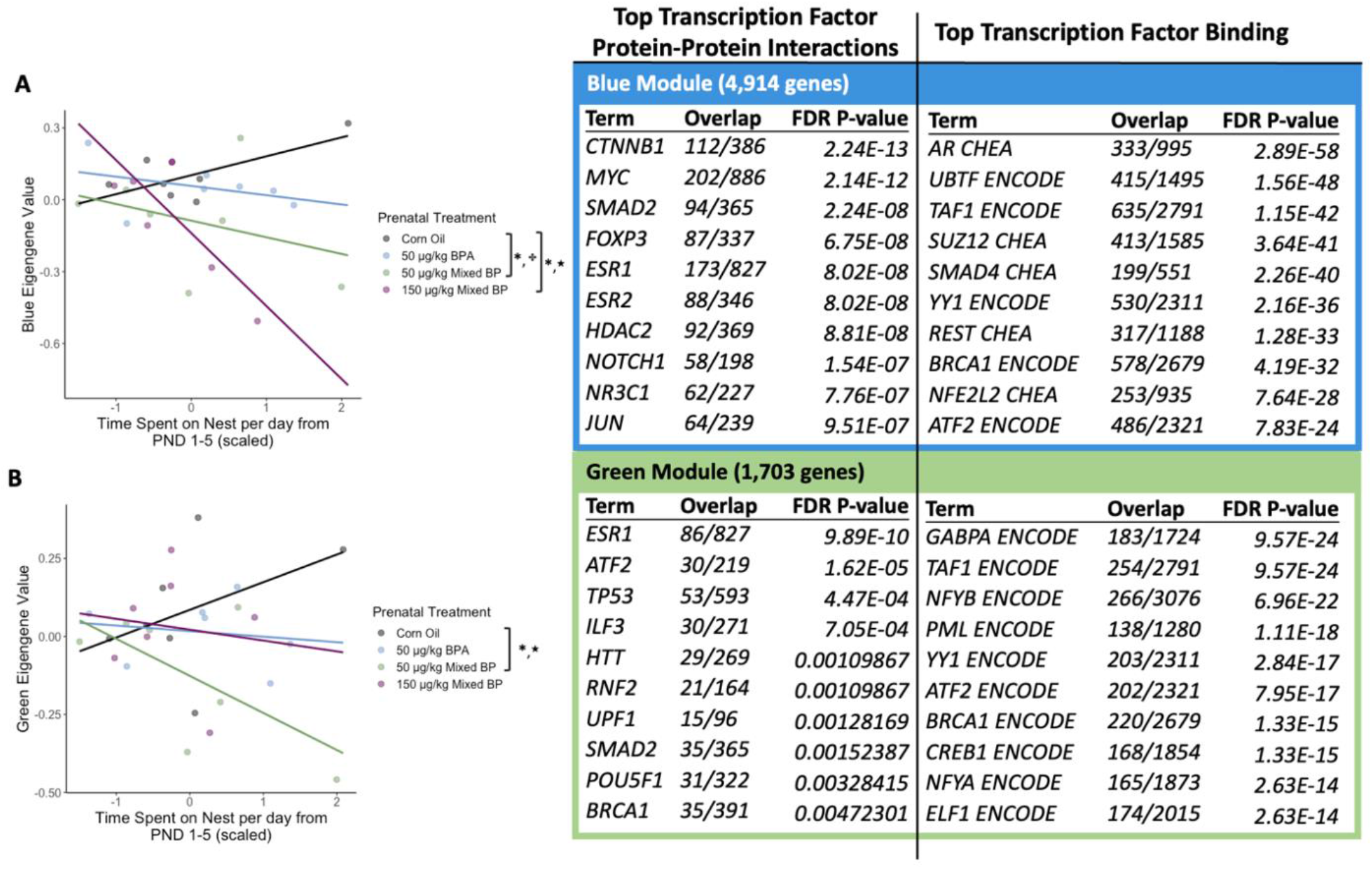
The interactive effects of prenatal bisphenol exposure and postnatal maternal care on transcriptomic profiles in the male amygdala. There were significant main effects or interactions with prenatal treatment and postnatal maternal care for the (A) blue module and (B) green module. Scatterplots are displayed with linear regression lines for each prenatal treatment group. * p < 0.05 main effect of prenatal treatment; ★ p < 0.05 interaction between prenatal treatment and postnatal maternal care; ✤ p < 0.10 interaction between prenatal treatment and postnatal maternal care

For the green module, there was a significant interaction between the 50 μg/kg Mixed BP exposure group and nest attendance (t = -2.118, p = 0.0468) compared to the Corn Oil group (Figure 7B). The top GO terms for the genes in the green module were RNA Splicing via Spliceosome, Substantia Nigra Development, mRNA Processing, RNA Splicing via Transesterification Reactions with Bulged Adenosine as Nucleophile, and Negative Regulation of Calcium Ion Transmembrane Transport. The top enriched transcription factors involved in protein-protein interactions with genes in the green module were ESR1, ATF2, TP53, ILF3, and HTT (Figure 7B). The top enriched transcription factors involved in facilitating transcription of genes in the green module were GABPA, TAF1, NFYB, PML, and YY1 (Figure 7B).

##### 3.4.2.2 Female Offspring

There were significant interactions between the 50 μg/kg Mixed BP exposure group and postnatal maternal care (Table 2). There was one DEG with a negative R^2^ change for the 50 μg/kg Mixed BP exposure group and nest attendance compared to the Corn Oil group, which included Hmga1-ps2 (Supplementary Figure 8A; FDR < 0.05). In addition, there was one DEG with a negative R^2^ change for the 50 μg/kg Mixed BP exposure group and licking/grooming compared to the Corn Oil group, which included Hmga1-ps2 (Supplementary Figure 8B; FDR < 0.05).

A total of eight eigengene modules were examined for differences between prenatal treatments and postnatal maternal care and their interactions. There were significant main effects or interactions found with two eigengene modules (Black and Yellow).

For the black module, there was a significant interaction between the 150 μg/kg Mixed BP exposure group and licking/grooming (t = 2.183, p = 0.0411) compared to the Corn Oil group (Figure 8A). The top GO terms for the genes in the black module were Translation, Macromolecule Biosynthetic Process, Peptide Biosynthetic Process, Cytoplasmic Translation, Proteasomal Protein Catabolic Process, and Ubiquitin-Dependent Protein Catabolic Process. The top enriched transcription factors involved in protein-protein interactions with genes in the black module were ESR1, ILF3, POLR2A, MYC, and FOXP3 (Figure 8A). The top enriched transcription factors involved in facilitating transcription of genes in the black module were TAF1, BRCA1, GABPA, CREB1, and YY1 (Figure 8A).

**Figure 8.**
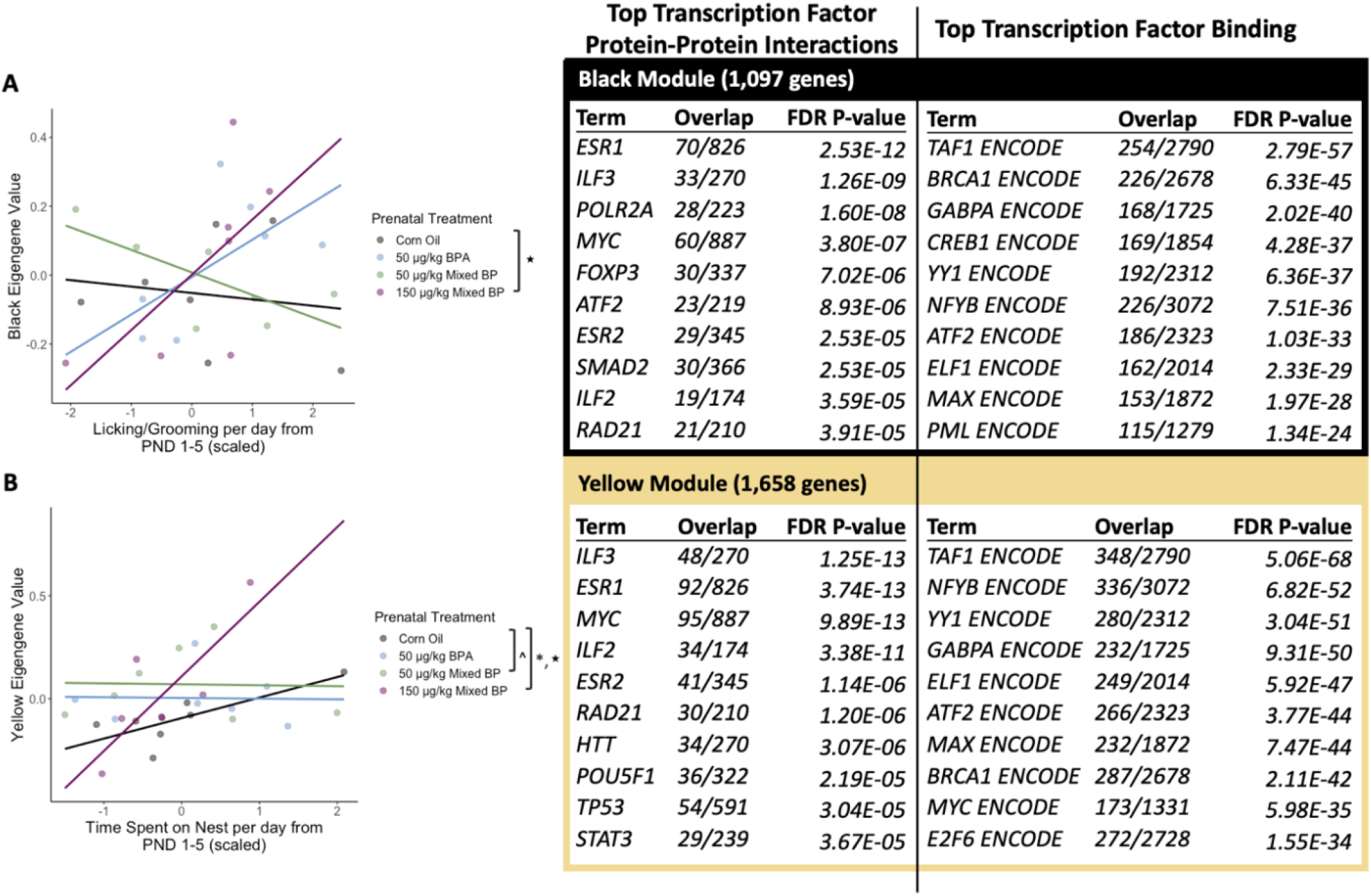
The interactive effects of prenatal bisphenol exposure and postnatal maternal care on transcriptomic profiles in the female amygdala. There were significant main effects or interactions with prenatal treatment and postnatal maternal care for the (A) black module and (B) yellow module. Scatterplots are displayed with linear regression lines for each prenatal treatment group. ^ p < 0.10 main effect of prenatal treatment; * p < 0.05 main effect of prenatal treatment; ★ p < 0.05 interaction between prenatal treatment and postnatal maternal care

For the yellow module, there was a significant interaction between the 150 μg/kg Mixed BP exposure group and licking/grooming (t = 2.205, p = 0.0393) compared to the Corn Oil group (Figure 8B). The top GO terms for the genes in the yellow module were Translation, Macromolecule Biosynthetic Process, Gene Expression, and Mitochondrial ATP Synthesis Coupled Electron Transport. The top enriched transcription factors involved in protein-protein interactions with genes in the yellow module were ILF3, ESR1, MYC, ILF2, and ESR2 (Figure 8B). The top enriched transcription factors involved in facilitating transcription of genes in the yellow module were TAF1, NFYB, YY1, GABPA, and ELF1 (Figure 8B).

## 4. Discussion

In this study, we examined the interactive effects of prenatal BP (BPA, BPF, BPS) exposure and postnatal maternal care on neurodevelopment and adult behavioral outcomes in rats. In addition, we investigated transcriptomic changes in the developing brain in response to prenatal BP exposure and postnatal maternal care. Overall, we determined that the effects of prenatal BP exposure on neurodevelopment and adult behavioral outcomes were dependent on postnatal maternal care, particularly variation in maternal care occurring during the first five days of life. We found significant or marginal moderation effects of postnatal maternal care on our outcome measures and the main effects of prenatal BP exposure were not apparent unless we accounted for postnatal maternal care in our statistical models. Notably, high levels of maternal care might also attenuate the effects of prenatal BP exposure on eye opening timing and adult attentional set shifting performance. In addition, we also found significant interactions between prenatal BP exposure and postnatal maternal care on differential gene expression in the developing medial prefrontal cortex and amygdala. Interestingly, a portion of the gene sets identified with WCGNA that were affected by prenatal BP exposure, postnatal maternal care, and/or their interactions were related to gene expression and translation processes and had prominent predicted interactions with estrogen receptor alpha (ESR1) and estrogen receptor beta (ESR2). Finally, we found these effects were dose-dependent and sex-specific, which have been reported in previous studies on prenatal BPA exposure (Arambula et al., 2018; Kubo et al., 2003; Kundakovic et al., 2013; Vandenberg, 2014; Vandenberg et al., 2013). Overall, these findings indicate that postnatal maternal care plays a critical role in the expression of the effects of prenatal BP exposure on neurodevelopment and adult behavior. These findings also suggest that changes in estrogen receptor signaling may be an important mechanism underlying the interactive effects of prenatal BP exposure and postnatal maternal care in the developing brain.

We hypothesized that the interactive effects between prenatal BP exposure and postnatal maternal care would influence our neurodevelopment and adult behavior outcome measures. We consistently found interactions between prenatal BP exposure and postnatal maternal care (measured by licking/grooming and nest attendance) on offspring outcomes and the main effects of prenatal BP exposure were dependent on postnatal maternal care. We used prevalence of eye opening at PND 15 as a neurodevelopmental milestone and found an overall reduction of eye-opening prevalence for offspring, particularly with males, with prenatal BP exposure. Higher maternal licking/grooming has been associated with acceleration of eye-opening time in rats (Ellenbroek et al., 2005; Uriarte et al., 2007) although prenatal BPA exposure has not been associated with changes in eye opening time (e.g., Ema et al., 2001; Nagao et al., 1999). One study with prenatal BPS exposure found a reduction in eye opening prevalence at PND 14 in mice, but the differences were not statistically significant (LaPlante et al., 2017). Our results also suggest that litters with high levels of licking/grooming have increased eye opening prevalence at PND 15 if they had prenatal BP exposure but decreased eye-opening prevalence at PND 15 for the control group. Given that both acceleration and deceleration of neurodevelopment are associated with adverse later-life psychiatric outcomes (Shaw et al., 2010), it is possible that higher maternal licking/grooming may be normalizing neurodevelopment instead of accelerating the pace of neurodevelopment in all contexts.

Our analyses indicate a reduction in adult attentional set shifting performance in offspring with prenatal BP exposure. These effects appear to be larger in female offspring than the male offspring and with prenatal BPA exposure than the BP mixture. We also found interactions between prenatal BP exposure and postnatal nest attendance on adult attentional set shifting performance. Prenatal BP exposure has been associated with impairments in executive function and cognition as well as increased diagnoses of attentional deficit/hyperactivity disorder in human populations (Bornehag et al., 2021; Braun et al., 2011, 2017; Rochester et al., 2018). In addition, higher maternal licking/grooming has been associated with improved attentional set shifting performance in rats (Lauby et al., 2019; Lovic & Fleming, 2004; McLean et al., 2010). In our study, nest attendance includes licking/grooming of pups but also other pup-directed behaviors such as nursing and thermotactile contact with the pups. The relationship between maternal behaviors other than licking/grooming and adult behavior has not been well-studied, and it is possible that multiple maternal behaviors could contribute to a single later-life outcome. Our results also suggest that female offspring with high levels of nest attendance have better performance in the attentional set shifting task if they had prenatal BP exposure but worse performance in the attentional set shifting task for the control group. The attentional set shifting task primarily assesses behavioral flexibility, but the neural mechanisms underlying high behavioral flexibility may conflict with selective, focused attention on a task (Aston-Jones et al., 1999). Therefore, it is possible that higher maternal care may also be normalizing performance on this task instead of improving performance in all contexts.

We found that male offspring with prenatal BP exposure, primarily the 50 μg/kg doses, have decreased anxiety-like behavior in the open field test. We also found interactions between prenatal BP exposure and postnatal nest attendance on adult anxiety-like behavior in the open field. Previous work in human populations have found prenatal BPA exposure is associated with impairments in emotional regulation at childhood (Braun et al., 2009, 2011; Perera et al., 2012, 2016; Roen et al., 2015). A previous study in our lab with prenatal BPA exposure in mice also found reductions in anxiety-like behavior, specifically in male offspring (Kundakovic et al., 2013). In addition, higher maternal licking/grooming has been associated with decreased anxiety-like behavior in rats (Caldji et al., 2000; Francis & Meaney, 1999). The post-hoc comparisons from the moderation analysis suggest that male offspring with prenatal BP exposure have decreased anxiety-like behavior if they received low levels of nest attendance but increased anxiety-like behavior if they received high levels of nest attendance. This is likely due to stronger effects of postnatal nest attendance on anxiety-like behavior in the control group compared to the prenatal BP exposure groups. These findings suggest that higher postnatal maternal care does not improve all later-life outcomes for offspring with prenatal BP exposure. Understanding the biological mechanisms underlying the interactive effects of prenatal BP exposure and postnatal maternal care might allow us to predict which later-life outcomes may be attenuated by higher levels of postnatal maternal care.

Finally, we found numerous differentially expressed genes (DEGs) and differentially expressed eigengene modules associated with interactions between prenatal BP exposure and postnatal maternal care. We identified more DEGs and differentially expressed eigengene modules for offspring with the prenatal BP mixture (both 50 μg/kg and 150 μg/kg) than the BPA-only group. In addition, we identified more DEGs and differentially expressed eigengene modules for postnatal nest attendance than licking/grooming. Highlighted DEGs for the male medial prefrontal cortex included Nrgn, Tuba4a, Nefl, and Mt3. Neurogranin (Nrgn) is linked to learning and memory in mice and higher levels of Nrgn are linked to synaptic dysfunction (Yin et al., 2013). Tubulin alpha 4a (Tuba4a) is associated with the development of post-mitotic neurons (Leca et al., 2023). Increases in neurofilament light polypeptide (Nefl) in serum are used a biomarker for axonal damage (Ceylan et al., 2023). Metallothionein 3 (Mt3) maintains copper and zinc levels in neurons and regulates the size of neurons as a growth inhibitor (Koh & Lee, 2020). Most of these DEGs reappear in the yellow eigengene module, which include biological processes related to ATP synthesis, oxidative phosphorylation, and neurotransmission. Highlighted DEGs for the female medial prefrontal cortex included Glra3. Glycine receptor alpha 3 (Glra3) is a subunit of the glycine receptor, which is a ubiquitously expressed inhibitory receptor in the brain. One previous study with prenatal synthetic glucocorticoid exposure also found changes in gene expression of Glra3 in the developing prefrontal cortex of guinea pigs (Constantinof et al., 2019). Highlighted DEGs for the male amygdala included Tafa2, Grip1, Robo1, and Epha3. TAFA chemokine like family member 2 (Tafa2) is associated with learning and memory as well as anxiety-like behavior in mice (Wang et al., 2018). Glutamate receptor interacting protein 1 (Grip1) is important for long term potentiation and learning and memory in mice (Tan et al., 2020). Roundabout guidance receptor 1 (Robo1) is involved in axon guidance and is associated with developmental dyslexia in human populations (Hannula-Jouppi et al., 2005). Ephrin tyrosine kinase a3 (Epha3) is also involved in axon guidance in mice (Nishikimi et al., 2011). These DEGs reappear in the blue eigengene module, which include biological processes related to neurotransmission, gene regulation, and neurodevelopment. Highlighted DEGs for the female amygdala included Hmga1-ps2. This is a pseudogene that is related to high mobility group AT-hook 1 (Hmga1) and the biological function of this gene is unknown.

Based on the GO terms for the differentially expressed eigengene modules in our analyses, genes related to gene transcription and protein translation and synthesis are altered by prenatal BP exposure and postnatal maternal care and could lead to widespread gene expression changes later in development. There were GO terms related to neurotransmission and neurodevelopment that were present in the male offspring but absent in the female offspring, which corresponds to the sex-specific effects with prenatal BP exposure and postnatal maternal care on our later-life outcome measures. This could be related to the high enrichment of androgen receptor (AR), which can be also altered by BPA exposure (Huang et al., 2019), as a transcription factor facilitating transcription of genes in the blue eigengene module in the male amygdala. However, a previous RNA-seq study with PND 1 amygdala found that prenatal BPA exposure was associated with DEGs related to synaptic plasticity and neurodevelopment in female offspring only (Arambula et al., 2018). The BP mixture used, inclusion of postnatal maternal care as an independent variable, and postnatal age of the pups in our study might account for those discrepancies. Finally, we do not find substantial or consistent enrichment of the estrogen receptors (ESR1 and ESR2) as transcription factors that facilitate transcription of genes in our eigengene modules. Instead, the genes in our eigengene modules have known interactions with ESR1 as well as ESR2 to a smaller extent. Although bisphenols are known to disrupt estrogen receptor signaling and estrogen receptors can modify gene transcription by binding to estrogen responsive elements in the DNA, estrogen receptors do not act alone in modifying gene transcription. In addition, estrogen receptors might be facilitating gene expression of other transcription factors during the perinatal period, so it is possible those transcription factors are showing higher enrichment by PND 10. Interestingly, the activity of ESR1 specifically can be enhanced or repressed by CREB1, BRCA1, and RAD1 (Berto et al., 2018; Lo et al., 2008; Prenzel et al., 2012) as well as induce expression of MYC and HTT (Nuzzo et al., 2016; Wang et al., 2011). CREB1 activity is also responsive to estradiol and BPA exposure (Quesada et al., 2002). These transcription factors consistently show high enrichment as transcription factors that facilitate gene expression and/or interact with the genes in our eigengene modules. This suggests that the interactive effects of prenatal BP exposure and postnatal maternal care might converge on estrogen receptor signaling changes. Higher postnatal maternal licking/grooming has been associated with increased Esr1 expression and responsiveness to estradiol (Champagne et al., 2003, 2006) but other aspects of estrogen receptor signaling have not been examined. In addition, we found other transcription factors that were repeatedly enriched in our eigengene modules, including TAF1, YY1, ILF3, and ATF2. Prenatal BPA exposure has been previously predicted to alter YY1 activity in the neonatal rat prefrontal cortex (Kanlayaprasit et al., 2021). However, these *in silico* predictions need to be tested experimentally in future work. It will also be important to understand the longitudinal changes in the interactions between ESR1 and other transcription factors and the long-term interactive effects of prenatal BP exposure and postnatal maternal care on gene regulation, including epigenetic mechanisms such as DNA methylation (Champagne & Curley, 2008; Kundakovic et al., 2013; Kundakovic & Champagne, 2011).

Previous work in rats and mice have found significant effects of gestational BPA or BPS exposure on maternal care provisioning (Catanese & Vandenberg, 2017; Kundakovic et al., 2013; Seta et al., 2005). However, in the current study, we did not observe these effects, likely due to the limited assessment of postnatal care which was restricted to one hour per day. Another caveat of this study is that the main effects and interactive effects of postnatal maternal care are correlative because they derive from natural observations. A similar limitation is that we do not have any control over the range of maternal care provided to offspring in our study and may be difficult to replicate across studies. To infer causal mechanisms, we would need to directly manipulate maternal care provided to the offspring and this will be a focus for future work. Finally, we used a limited range of doses for the BP treatment groups so we cannot fully assess the nonmonotonic response curve seen in other studies with prenatal BPA exposure. A wider range of doses as well as smaller doses than the ones chosen for our study will be important to understand the effects of environmentally relevant mixtures of bisphenols on human health.

Overall, our study suggests that postnatal maternal care has a substantial influence on neurodevelopment and behavior for offspring with prenatal BP exposure. Importantly, high postnatal maternal might be a protective factor against the adverse effects of prenatal BP exposure on some later-life outcomes. Our study also showed that the main effects of prenatal BP exposure were masked unless we accounted for maternal care. This emphasizes the need to consider multiple co-occurring exposures when assessing the effects of one specific exposure on phenotype. Finally, our analysis on gene expression changes in two relevant brain areas in male and female offspring suggest that changes in estrogen receptor signaling at different levels of gene regulation may be one possible mechanism in which prenatal BP exposure and postnatal maternal care exert their effects. Understanding the underlying biological mechanisms involved might allow us to identify potential avenues to mitigate the adverse effects of prenatal BP exposure and improve health and well-being in human populations.

## Supporting information

Supplementary Methods and Results

## Acknowledgements

We thank Megan Nguyen, Isha Agarwal, Taylor Hite, and Madeline Severson for their assistance in the animal experiments. We also thank Dr. Dennis Wylie for his assistance in conducting and troubleshooting the tag-seq bioinformatic analyses. This research was supported by a R01 grant (1R01ES030950) from the National Institute for Environmental Health Sciences (NIEHS) to A.E.M. and F.A.C. and a F32 postdoctoral fellowship (1F32ES035254) from the NIEHS to S.C.L.

