## Supplementary Figures for "Postnatal maternal care moderates the effects of prenatal bisphenol exposure on offspring neurodevelopmental, behavioral, and transcriptomic outcomes"

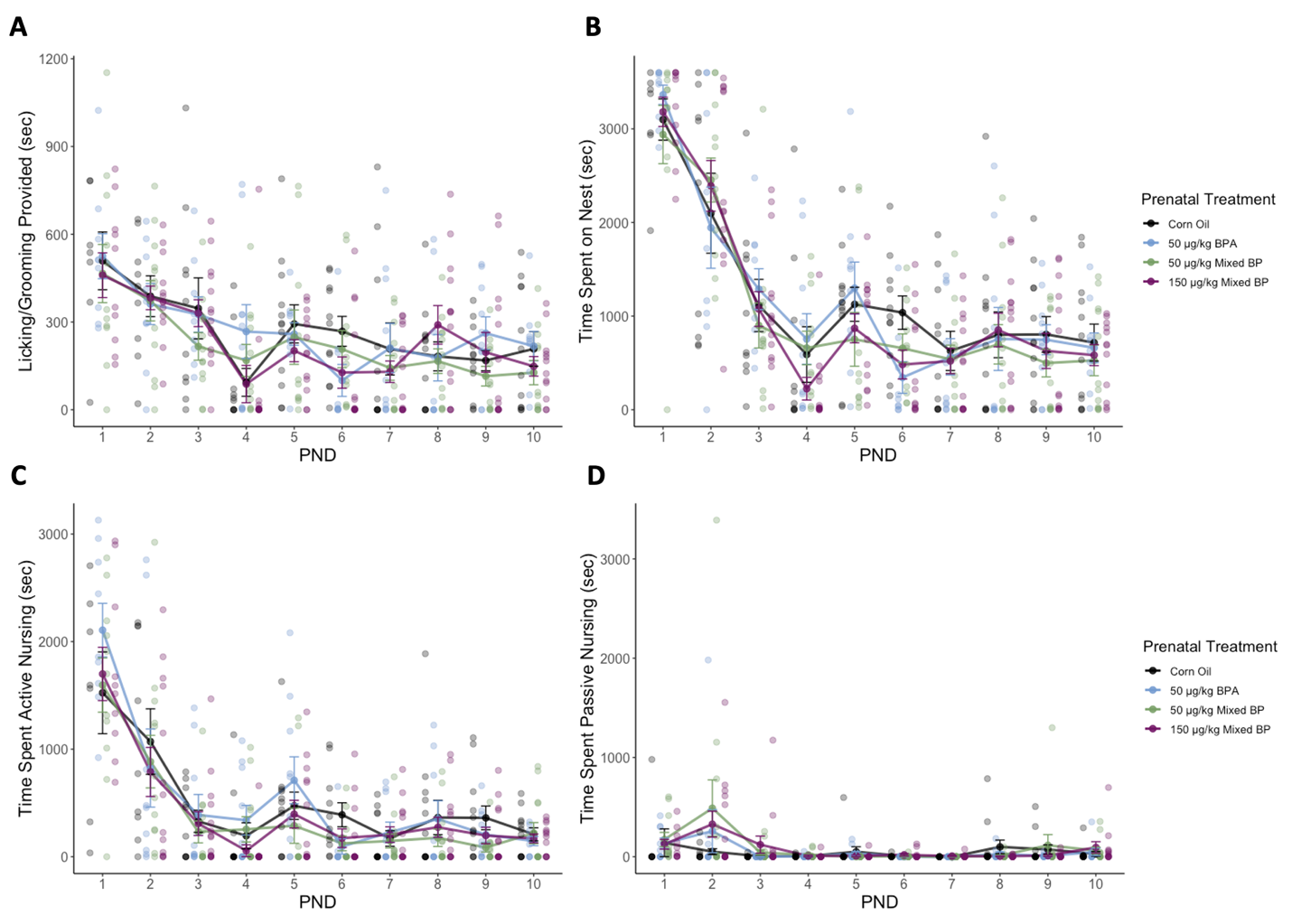

**Supplementary Figure 1.** There was no effect of gestational bisphenol exposure on postnatal maternal care provisioning. No main effects of prenatal treatment were found with (A) licking/grooming provisioning, (B) nest attendance, (C) active nursing, or (D) passive nursing. Line graphs are displayed with mean +/- SEM for each prenatal treatment group with individual datapoints.

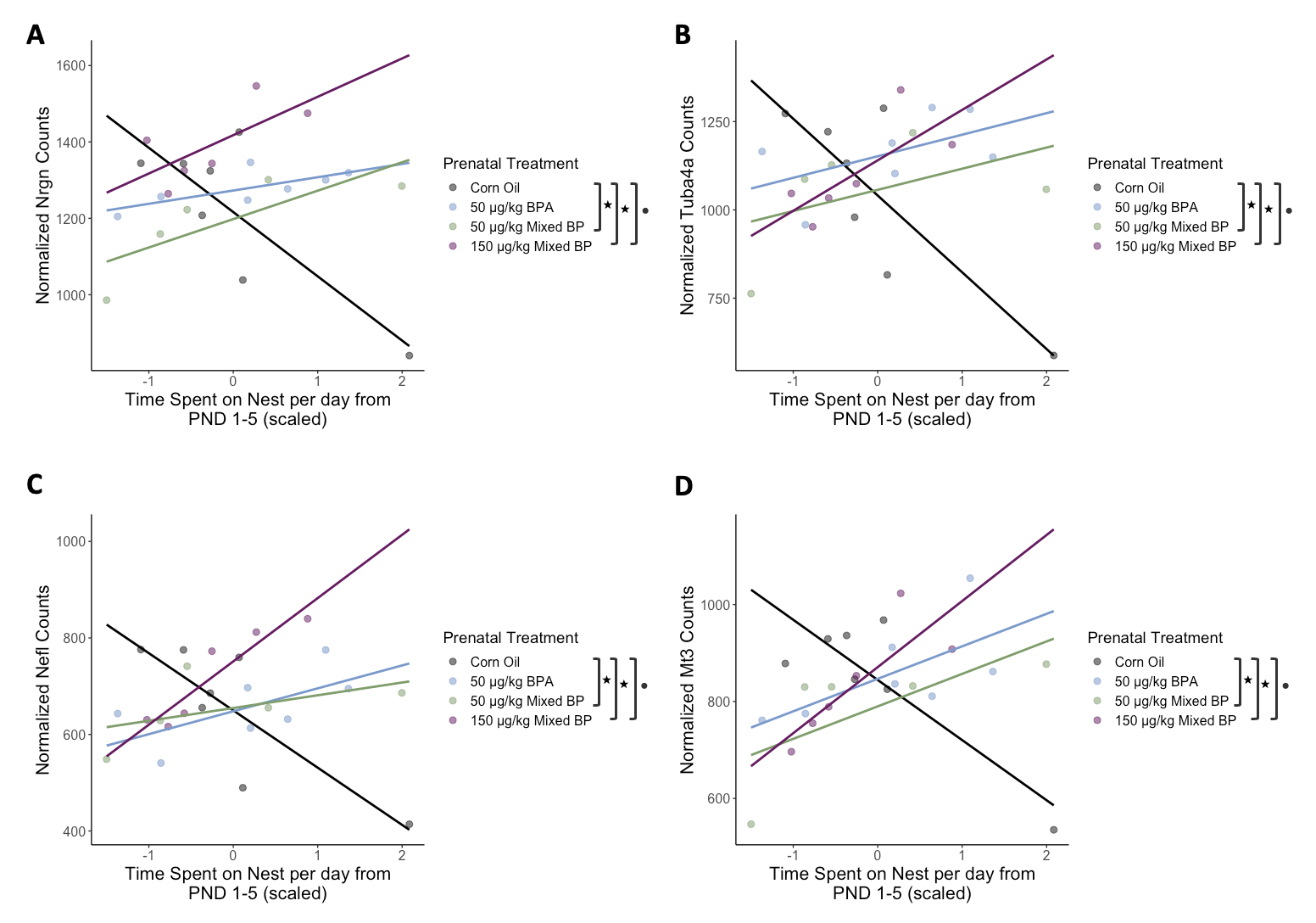

**Supplementary Figure 2.** The interactive effects of prenatal bisphenol exposure and postnatal maternal care on select differentially expressed genes in the male medial prefrontal cortex. A significant main effect of postnatal nest attendance and significant interactions between prenatal treatment (50 μg/kg Mixed BP and 150 μg/kg Mixed BP) and postnatal nest attendance were found for (A) Nrgn, (B) Tuba4a, (C) Nefl, and (D) Mt3. Scatterplots are displayed with linear regression lines for each prenatal treatment group. ● p < 0.05 main effect of postnatal maternal care; + p < 0.10 main effect of postnatal maternal care; ★ p < 0.05 interaction between prenatal treatment and postnatal maternal care

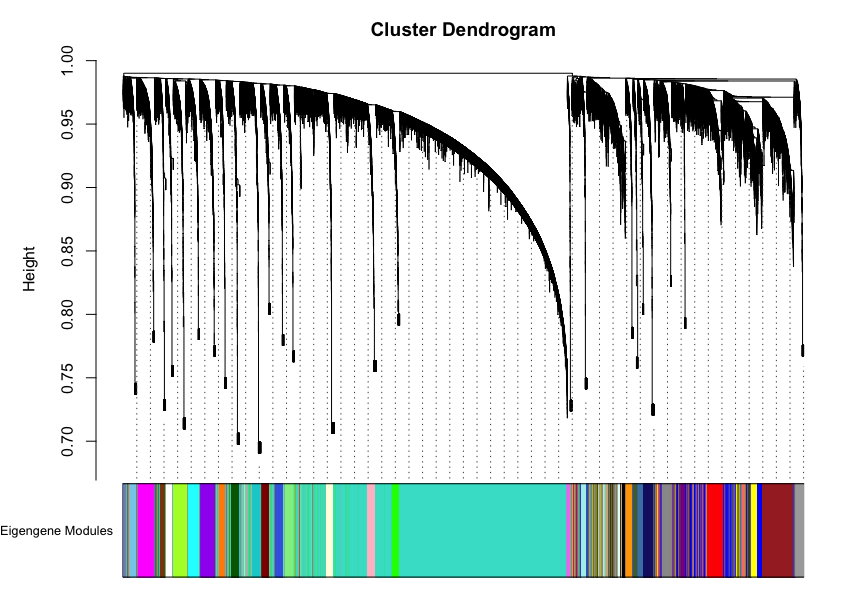

**Supplementary Figure 3.** Weighted gene co-expression network analysis identified 35 eigengene modules for the male medial prefrontal cortex.

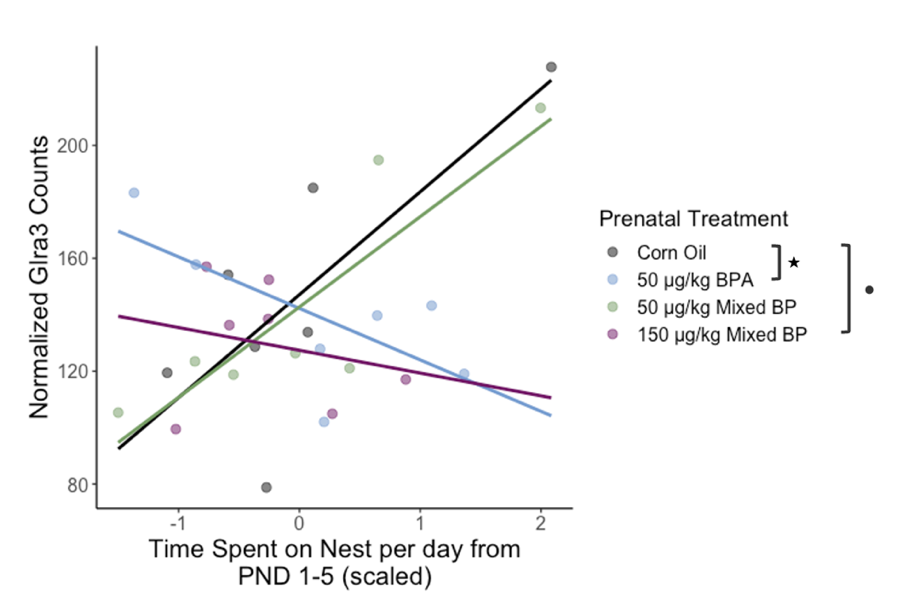

**Supplementary Figure 4.** The interactive effects of prenatal bisphenol exposure and postnatal maternal care on Glra3 gene expression in the female medial prefrontal cortex. There was a significant main effect of postnatal nest attendance and significant interaction between prenatal treatment (50 μg/kg Mixed BP) and postnatal nest attendance. Scatterplot is displayed with linear regression lines for each prenatal treatment group. ● p < 0.05 main effect of postnatal maternal care; ★ p < 0.05 interaction between prenatal treatment and postnatal maternal care

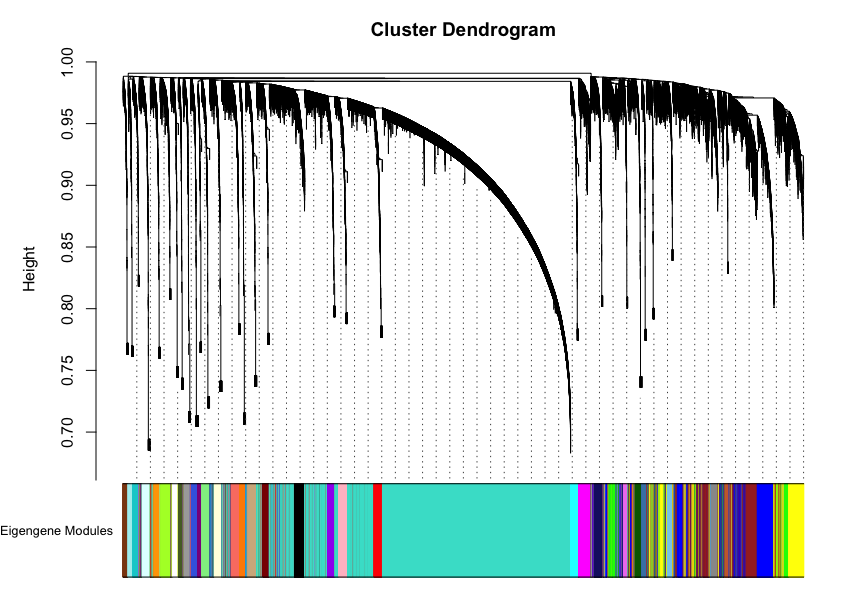

**Supplementary Figure 5.** Weighted gene co-expression network analysis identified 37 eigengene modules for the female medial prefrontal cortex.

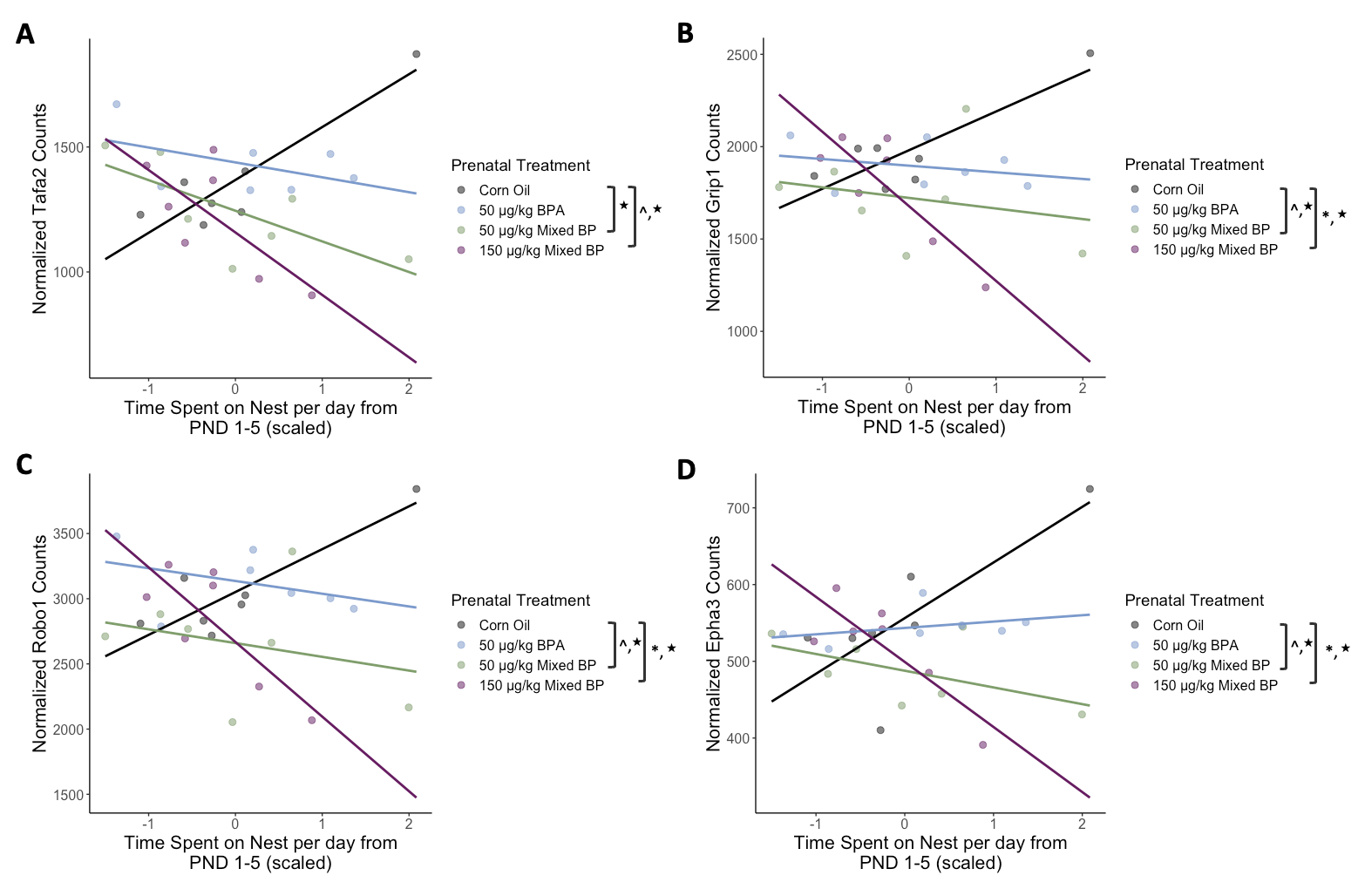

**Supplementary Figure 6.** The interactive effects of prenatal bisphenol exposure and postnatal maternal care on select differentially expressed genes in the male amygdala. Significant or marginal main effects of prenatal treatment (50 μg/kg Mixed BP and 150 μg/kg Mixed BP) and significant interactions between prenatal treatment and postnatal nest attendance were found for (A) Tafa2, (B) Grip1, (C) Robo1, and (D) Epha3. Scatterplots are displayed with linear regression lines for each prenatal treatment group. ^ p < 0.10 main effect of prenatal treatment; * p < 0.05 main effect of prenatal treatment; ★ p < 0.05 interaction between prenatal treatment and postnatal maternal care

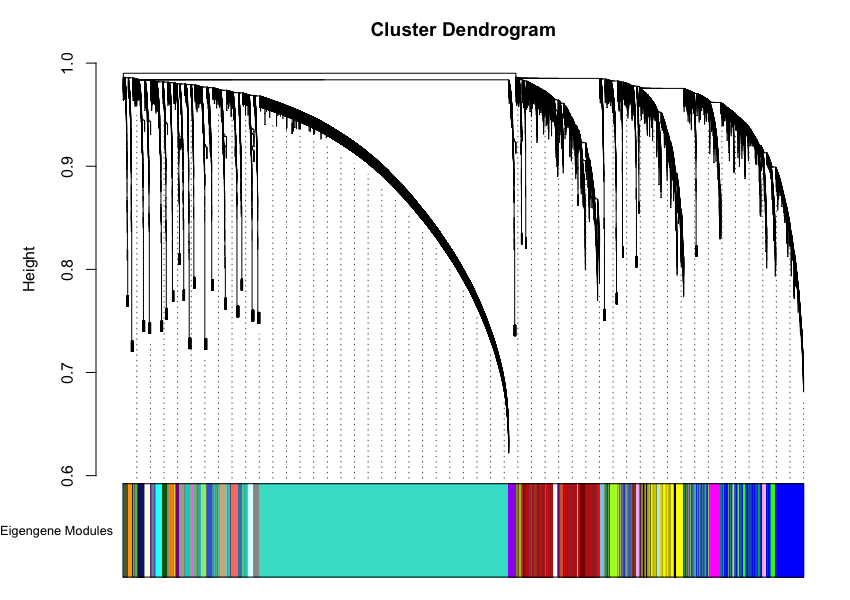

**Supplementary Figure 7.** Weighted gene co-expression network analysis identified 42 eigengene modules for the male amygdala.

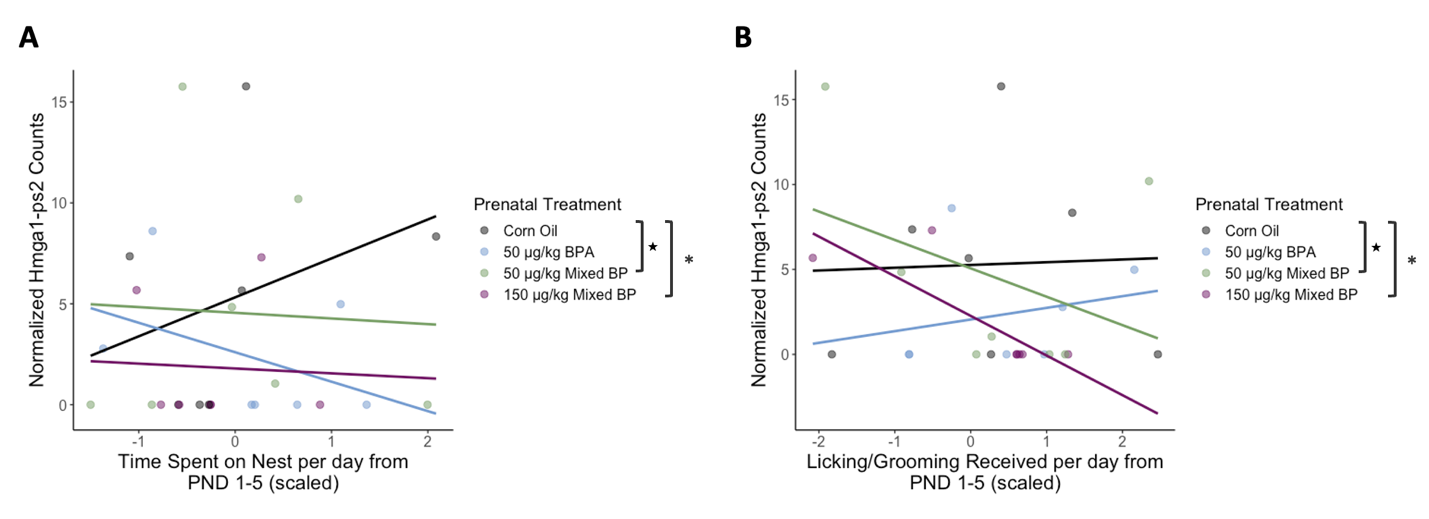

**Supplementary Figure 8.** The interactive effects of prenatal bisphenol exposure and postnatal maternal care on Hmga1-ps2 gene expression in the female amygdala. There was a significant main effect of prenatal treatment (150 μg/kg Mixed BP) and significant interactions between (A) prenatal treatment (50 μg/kg Mixed BP) and postnatal nest attendance and (B) prenatal treatment (50 μg/kg Mixed BP) and licking/grooming. Scatterplots are displayed with linear regression lines for each prenatal treatment group. * p < 0.05 main effect of prenatal treatment; ★ p < 0.05 interaction between prenatal treatment and postnatal maternal care

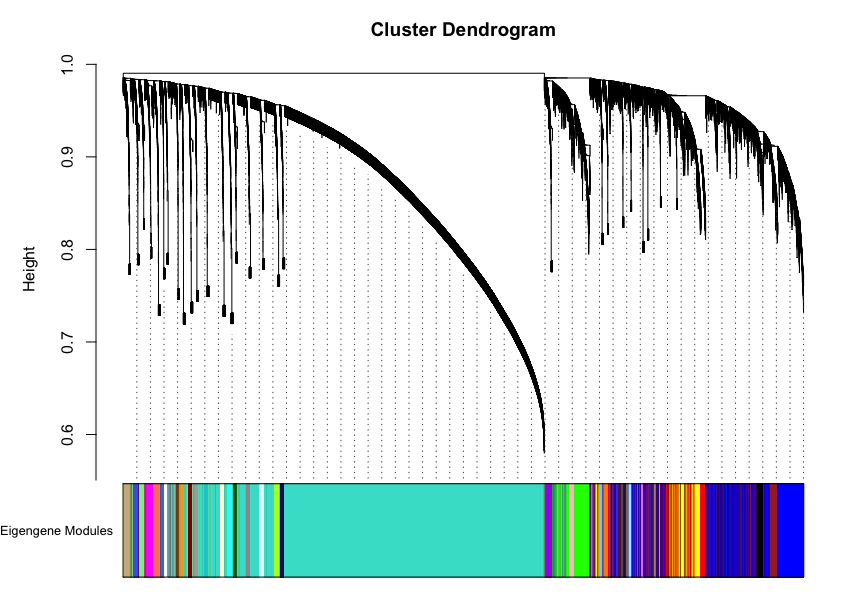

**Supplementary Figure 9.** Weighted gene co-expression network analysis identified 36 eigengene modules in the female amygdala.
