## Supplementary Methods and Results for "Postnatal maternal care moderates the effects of prenatal bisphenol exposure on offspring neurodevelopmental, behavioral, and transcriptomic outcomes"

**Supplementary Materials and Methods**

**Statistical Analysis**

All statistical analyses were performed using R version 4.1.3 (R Core Team, 2023). Linear mixed models were performed using the lme4 R package and F-tests were performed using the ANOVA (Type III) function in the cars R package. Significant main effects of prenatal treatment were followed with t-tests between the Corn Oil group and each of the bisphenol treatment groups using the lmerTest R package. Batch of breeding was used as a random factor.

To assess if prenatal bisphenol exposure significantly influences postnatal maternal care provisioning, a linear mixed model was used with prenatal treatment and postnatal day as independent variables.

**Supplementary Results**

**Effects of Prenatal Bisphenol Exposure on Postnatal Maternal Care Provisioning**

There were no main effects of prenatal treatment or interaction between prenatal treatment and postnatal day (all p’s > 0.10) on postnatal licking/grooming (Supplementary Figure 1A), nest attendance (Supplementary Figure 1B), active nursing (Supplementary Figure 1C), or passive nursing (Supplementary Figure 1D).

**Postnatal Maternal Care Moderates the Effects of Prenatal Bisphenol Exposure on Neurodevelopment**

Moderation analyses with PROCESS confirmed the marginal interactions between prenatal treatment and postnatal licking/grooming received for total pups in the litter with eyes open (R^2^ change = 0.1636, F_3,37_ = 2.5828, p = 0.06790) and male pups in the litter with eyes open (R^2^ change = 0.1681, F_3,37_ = 2.8123, p = 0.0526). For offspring with low levels of licking/grooming received, there was a marginal reduction in prevalence in eye-opening between the Corn Oil group and the 50 μg/kg BPA (t = -1.8343, p = 0.0747) exposure group and a significant reduction between the Corn Oil group and the 50 μg/kg Mixed BP (t = -2.7486, p = 0.0092), and 150 μg/kg Mixed BP (t = -2.4418, p = 0.0195) exposure groups (Table 1). For offspring with mid levels of licking/grooming received, there was a marginal reduction in prevalence in eye-opening between the Corn Oil group and the 50 μg/kg Mixed BP (t = -1.8337, p = 0.0748) exposure group. There were no significant differences between prenatal treatment groups for offspring with high levels of licking/grooming received. Similarly, for male offspring with low levels of licking/grooming received, there was a marginal reduction in prevalence in eye-opening with the 50 μg/kg BPA (t = -1.6980, p = 0.0979) exposure group compared to the Corn Oil group and a significant reduction with the 50 μg/kg Mixed BP (t = -3.4311, p = 0.0015), and 150 μg/kg Mixed BP (t = -2.5692, p = 0.0144) exposure groups compared to the Corn Oil group (Table 1). For male offspring with mid levels of licking/grooming received, there was a significant reduction in prevalence in eye-opening with the 50 μg/kg Mixed BP (t = -2.2922, p = 0.0277) exposure group compared to the Corn Oil group (Table 1). There were no significant differences between prenatal treatment groups for male offspring with high levels of licking/grooming received (Table 1).

**Postnatal Maternal Care Moderates the Effects of Prenatal Bisphenol Exposure on Adult Behavior**

**Attentional Set-Shifting Performance**

Moderation analyses with PROCESS confirmed the marginal interaction between prenatal treatment and postnatal nest attendance for percent correct trials in male offspring (R^2^ change = 0.1725, F_3,35_ = 2.8063, p = 0.0539) and the significant interaction between prenatal treatment and postnatal nest attendance for trials to reach criterion in female offspring (R^2^ change = 0.2308, F_3,32_ = 3.3530, p = 0.0310). For male offspring with low levels of nest attendance, there was a marginal reduction in percent correct trials with the 50 μg/kg BPA (t = -1.8245, p = 0.0766) exposure group compared to the Corn Oil group (Table 1). There were no significant differences between prenatal treatment groups for male offspring with mid or high levels of nest attendance (Table 1). For female offspring with low levels of nest attendance, there was a significant increase in the number of trials to reach criterion with the 150 μg/kg Mixed BP (t = 2.2235, p = 0.0334) exposure group compared to the Corn Oil group. For female offspring with high levels of nest attendance, there was a marginal reduction in the number of trials to reach criterion with the 150 μg/kg Mixed BP (t = -1.7268, p = 0.0938) exposure group compared to the Corn Oil group and a significant reduction in the number of trials to reach criterion with the 50 μg/kg BPA (t = -2.0609, p = 0.0475) exposure group compared to the Corn Oil group (Table 1). There were no significant differences between prenatal treatment groups for female offspring with mid levels of nest attendance (Table 1).

**Anxiety-Like Behavior**

Moderation analyses with PROCESS confirmed the significant interaction between prenatal treatment and postnatal nest attendance for time spent in the center of the open field for male offspring (R^2^ change = 0.2273, F_3,76_ = 8.1170, p = < 0.001). For male offspring with low levels of nest attendance, there was a marginal increase in time spent in the center of the open field with the 50 μg/kg BPA (t = 1.7897, p = 0.0775) exposure group compared to the Corn Oil group and a significant increase with the 50 μg/kg Mixed BP (t = 2.4623, p = 0.0161) exposure group compared to the Corn Oil group (Table 1). For male offspring with mid levels of nest attendance, there was a marginal reduction in time spent in the center of the open field with the 50 μg/kg Mixed BP (t = -1.8663, p = 0.0775) exposure group compared to the Corn Oil group and a significant reduction with the 150 μg/kg Mixed BP (t = -2.0481, p = 0.044) exposure group compared to the Corn Oil group (Table 1). For male offspring with high levels of nest attendance, there was a significant reduction in time spent in the center of the open field with the 50 μg/kg BPA (t = -3.2814, p = 0.0016), 50 μg/kg Mixed BP (t = -4.4173, p = < 0.001), and 150 μg/kg Mixed BP (t = -2.5806, p = 0.0118) exposure groups compared to the Corn Oil group (Table 1).

**Interactive Effects of Prenatal Bisphenol Exposure and Postnatal Maternal Care on PND 10 Brain Gene Expression**

**Medial Prefrontal Cortex**

**Male Offspring**

A total of 23,155 genes were analyzed for differential expression in the male medial prefrontal cortex. There were two downregulated DEGs found in the 150 μg/kg Mixed BP exposure group compared to the Corn Oil group, which included Hmga1-ps2 and LOC120099710 (Table 3; all FDR < 0.05). In addition, there were 25 negatively associated DEGs found with postnatal nest attendance, where male offspring with higher nest attendance had lower expression of these genes (Table 3). The top negatively associated DEGs included Nrgn, Tuba4a, and Nefl (all FDR < 0.05; Supplementary Table 1). Finally, there were significant interactions between the 50 μg/kg Mixed BP and 150 μg/kg Mixed BP exposure groups and postnatal nest attendance (Table 3). There were 24 DEGs with a positive R^2^ change for the 50 μg/kg Mixed BP exposure group and nest attendance compared to the Corn Oil group (Supplementary Table 2). There were 22 DEGs with a positive R^2^ change and 3 DEGS with a negative R^2^ change for the 150 μg/kg Mixed BP exposure group and nest attendance compared to the Corn Oil group (Supplementary Table 3). Some common DEGs with a positive R^2^ change included Nrgn (Supplementary Figure 2A), Tuba4a (Supplementary Figure 2B), Nefl (Supplementary Figure 2C), and Mt3 (Supplementary Figure 2D) (all FDR < 0.05).

Count data from 35,310 genes were used to construct networks for the WCGNA and were clustered into 35 eigengene modules (Supplementary Figure 3). A total of 10 eigengene modules were examined for differences between prenatal treatments and postnatal maternal care and their interactions. There were significant main effects or interactions found with five eigengene modules (Black, Brown, Lightcyan, Tan, and Yellow).

For the black module (containing 947 genes), there was a significant main effect of postnatal nest attendance (t = -3.017, p = 0.00777; Figure 5A). There were also significant interactions between the 50 μg/kg BPA exposure group and nest attendance (t = 2.170, p = 0.04443), 50 μg/kg Mixed BP exposure group and nest attendance (t = 2.662, p = 0.01641), and the 150 μg/kg Mixed BP exposure group and nest attendance (t = 2.242, p = 0.03861) compared to the Corn Oil group (Figure 5A). The top GO terms for the genes in the black module were Macromolecule Biosynthetic Process, Translation, Peptide Biosynthetic Process, Gene Expression, Regulation of Apoptotic Process, and Protein Transport. The top enriched transcription factors involved in protein-protein interactions with genes in the black module were MYC, ESR1, TP53, ILF3, and ATF2 (Figure 5A). The top enriched transcription factors involved in facilitating transcription of genes in the black module were GABPA, ELF1, TAF1, NFYB, and YY1 (Figure 5A).

For the brown module (containing 2,035 genes), there was a significant difference between the 150 μg/kg Mixed BP exposure group and Corn Oil group (t = 2.236, p = 0.0391; Figure 5B). There was a marginal main effect of postnatal licking/grooming (t = 1.885, p = 0.0766) and a significant interaction between the 50 μg/kg BPA exposure group and licking/grooming (t = -2.704, p = 0.0150) compared to the Corn Oil group (Figure 5B). The top GO terms for the genes in the brown module were Translation, Gene Expression, Cytoplasmic Translation, and Peptide Biosynthetic Process. The top enriched transcription factors involved in protein-protein interactions with genes in the brown module were ESR1, MYC, ESR2, ILF3, and RAD21 (Figure 5B). The top enriched transcription factors involved in facilitating transcription of genes in the brown module were TAF1, ATF2, YY1, BRCA1, and MYC (Figure 5B).

For the lightcyan module (containing 501 genes), there was a significant main effect of postnatal nest attendance (t = -3.257, p = 0.00464; Figure 5C). There were also significant interactions between the 50 μg/kg BPA exposure group and nest attendance (t = 2.418, p = 0.02714) and the 50 μg/kg Mixed BP exposure group and nest attendance (t = 2.174, p = 0.04415) compared to the Corn Oil group (Figure 5C). There were no significantly enriched GO terms or transcription factor protein-protein interactions for the genes in the lightcyan module. The top enriched transcription factors involved in facilitating transcription of genes in the lightcyan module were ATF2, ZBTB7A, and BRCA1 (Figure 5C).

For the tan module (containing 693 genes), there was a significant main effect of postnatal nest attendance (t = -2.848, p = 0.0111; Figure 5D). There was also a marginal interaction between the 50 μg/kg Mixed BP exposure group and nest attendance (t = 1.917, p = 0.0722) compared to the Corn Oil group (Figure 6I). There were no significantly enriched GO terms and the top transcription factor protein-protein interaction, ESR1, was marginally significant (FDR = 0.0702) for the genes in the tan module (Figure 5D). The top enriched transcription factors involved in facilitating transcription of genes in the tan module were CREB1, NFYB, NFYA, ZBTB7A, and PML (Figure 5D).

For the yellow module (containing 1,320 genes), there was a significant main effect of postnatal nest attendance (t = -2.527, p = 0.0217; Figure 5E). There was a significant interaction between the 50 μg/kg Mixed BP exposure group and nest attendance (t = 2.667, p = 0.0162) and marginal interactions between the 50 μg/kg BPA exposure group and nest attendance (t = 1.750, p = 0.0981) and the 150 μg/kg Mixed BP exposure group and nest attendance (t = 2.069, p = 0.0541) compared to the Corn Oil group (Figure 5E). The top GO terms for the genes in the yellow module were Proton Motive Force-Driven ATP Synthesis, Proton Motive Force-Driven Mitochondrial ATP Synthesis, Oxidative Phosphorylation, Signal Release from Synapse, and Neurotransmitter Secretion. The top enriched transcription factors involved in protein-protein interactions with genes in the yellow module were HTT, ESR1, ILF2, ILF3, and ATF2 (Figure 5E). The top enriched transcription factors involved in facilitating transcription of genes in the yellow module were TAF1, BRCA1, NRF1, ATF2, and YY1 (Figure 5E). The DEGs Nrgn, Tuba4a, and Nefl were found in this module.

**Female Offspring**

A total of 23,303 genes were analyzed for differential expression in the female medial prefrontal cortex. There was one downregulated DEG found in the 150 μg/kg Mixed BP exposure group compared to the Corn Oil group, which included Mir6332 (Table 3; FDR < 0.05). In addition, there was one negatively associated DEG found with postnatal nest attendance, where female offspring with higher nest attendance had lower expression of Glra3 (Supplementary Figure 4; FDR < 0.05). Finally, there were significant interactions between the 50 μg/kg BPA and 150 μg/kg Mixed BP exposure groups and postnatal nest attendance (Table 3). There was one DEG with a negative R^2^ change for the 50 μg/kg BPA exposure group and nest attendance compared to the Corn Oil group, which included Glra3 (Supplementary Figure 4; FDR < 0.05). There was one DEG with a negative R^2^ change for the 150 μg/kg BPA exposure group and nest attendance compared to the Corn Oil group, which included Dcx (FDR < 0.05). Glra3 was listed as the second top gene by unadjusted p-value but did not pass FDR p-value correction (FDR = 0.515).

Count data from 35,650 genes were used to construct networks for the WGCNA and were clustered into 37 eigengene modules (Supplementary Figure 5). A total of nine eigengene modules were examined for differences between prenatal treatments and postnatal maternal care and their interactions. There were significant main effects or interactions found with two eigengene modules (Brown and Turquoise).

For the brown module (containing 2,431 genes), there was a significant main effect of licking/grooming (t = 2.192, p = 0.04038; Figure 6A). There were also significant interactions between the 50 μg/kg BPA exposure group and licking/grooming (t = -2.141, p = 0.04478) and the 50 μg/kg Mixed BP exposure group and licking/grooming (t = -3.431, p = 0.00265) compared to the Corn Oil group (Figure 6A). The top GO terms for the genes in the brown module were Cytoplasmic Translation, Macromolecule Biosynthetic Process, Gene Expression, and Translation. The top enriched transcription factors involved in protein-protein interactions with genes in the brown module were ESR1, ILF3, RAD21, POU5F1, and MYC (Figure 6A). The top enriched transcription factors involved in facilitating transcription of genes in the brown module were TAF1, YY1, ATF2, BRCA1, and MYC (Figure 6A).

For the turquoise module (containing 14,733 genes), there were significant interactions between the 50 μg/kg BPA exposure group and licking/grooming (t = 2.102, p = 0.0484), the 50 μg/kg Mixed BP exposure group and licking/grooming (t = 2.758, p = 0.0121), and the 150 μg/kg Mixed BP exposure group and licking/grooming (t = 2.452, p = 0.0235) compared to the Corn Oil group (Figure 6B). There were no significantly enriched GO terms, transcription factor protein-protein interactions, or transcription factors that facilitate transcription of the genes in the turquoise module. The DEG Glra3 was found in this module.

**Amygdala**

**Male Offspring**

A total of 21,261 genes were analyzed for differential expression in the male amygdala. There were two upregulated DEGs and 285 downregulated DEGs found in the 50 μg/kg Mixed BP exposure group (Supplemental Table 4) and one upregulated DEG and 242 downregulated DEGs found in the 150 μg/kg Mixed BP exposure group (Supplemental Table 5) compared to the Corn Oil group (Table 3). Some common downregulated DEGs included Erbb4, Nlgn1, and Nrg3 (FDR < 0.10 for 50 μg/kg Mixed BP and FDR < 0.05 for 150 μg/kg Mixed BP). There were no DEGs associated with postnatal maternal care, but there were significant interactions between the 50 μg/kg Mixed BP and 150 μg/kg Mixed BP exposure groups and postnatal maternal care (Table 3). There were 17 DEGs with a positive R^2^ change and 223 DEGs with a negative R^2^ change for the 50 μg/kg Mixed BP exposure group and nest attendance compared to the Corn Oil group (Supplementary Table 6). There were 16 DEGs with a positive R^2^ change and 500 DEGs with a negative R^2^ change for the 150 μg/kg Mixed BP exposure group and nest attendance compared to the Corn Oil group (Supplementary Table 7). Some common DEGs with a negative R^2^ change included Tafa2 (Supplementary Figure 6A), Grip1 (Supplementary Figure 6B), Robo1 (Supplementary Figure 6C), and Epha3 (Supplementary Figure 6D; all FDR < 0.05). In addition, there was one DEG with a positive R^2^ change and one DEG with a negative R^2^ change for the 50 μg/kg Mixed BP exposure group and licking/grooming received compared to the Corn Oil group, which included Cdh13 and Col1a2 respectively (Table 3; all FDR < 0.10).

Count data from 36,473 genes were used to construct networks for the WCGNA and were clustered into 42 eigengene modules (Supplementary Figure 7). A total of 14 eigengene modules were examined for differences between prenatal treatments and postnatal maternal care and their interactions. There were significant main effects or interactions found with two eigengene modules (Blue and Green).

For the blue module (containing 4,914 genes), there were significant differences between the 50 μg/kg Mixed BP exposure group and Corn Oil group (t = -2.227, p = 0.03756) and the 150 μg/kg Mixed BP exposure group and Corn Oil group (t = -2.726, p = 0.01302; Figure 7A). There was also a significant interaction between the 150 μg/kg Mixed BP exposure group and nest attendance (t = -3.226, p = 0.00423) and a marginal interaction between the 50 μg/kg Mixed BP exposure group and nest attendance (t = -1.733, p = 0.09849) compared to the Corn Oil group (Figure 7A). The top GO terms for the genes in the blue module were Modulation of Chemical Synaptic Transmission, Chemical Synaptic Transmission, Nervous System Development, Regulation of Transcription by RNA Polymerase II, Anterograde Trans-Synaptic Signaling and Axonogenesis. The top enriched transcription factors involved in protein-protein interactions with genes in the blue module were CTNNB1, MYC, SMAD2, FOXP3, and ESR1 (Figure 7A). The top enriched transcription factors involved in facilitating transcription of genes in the blue module were AR, UBTF, TAF1, SUZ12, and SMAD4 (Figure 7A). The DEGs Erbb4, Nlgn1, Nrg3, Tafa2, Grip1, Robo1, Epha3, and Cdh13 were found in this module.

For the green module (containing 1,703 genes), there was a significant difference between the 50 μg/kg Mixed BP exposure group and Corn Oil group (t = -2.178, p = 0.0415; Figure 7B). There was also a significant interaction between the 50 μg/kg Mixed BP exposure group and nest attendance (t = -2.118, p = 0.0468) compared to the Corn Oil group (Figure 7B). The top GO terms for the genes in the green module were RNA Splicing via Spliceosome, Substantia Nigra Development, mRNA Processing, RNA Splicing via Transesterification Reactions with Bulged Adenosine as Nucleophile, and Negative Regulation of Calcium Ion Transmembrane Transport. The top enriched transcription factors involved in protein-protein interactions with genes in the green module were ESR1, ATF2, TP53, ILF3, and HTT (Figure 7B). The top enriched transcription factors involved in facilitating transcription of genes in the green module were GABPA, TAF1, NFYB, PML, and YY1 (Figure 7B).

**Female Offspring**

A total of 21,799 genes were analyzed for differential expression in the female amygdala. There was one downregulated DEG found in the 150 μg/kg Mixed BP exposure group compared to the Corn Oil group, which included Hmga1-ps2 (Table 3; FDR < 0.05). There were no DEGs associated with postnatal maternal care, but there were significant interactions between the 50 μg/kg Mixed BP exposure group and postnatal maternal care (Table 3). There was one DEG with a negative R^2^ change for the 50 μg/kg Mixed BP exposure group and nest attendance compared to the Corn Oil group, which included Hmga1-ps2 (Supplementary Figure 8A; FDR < 0.05). In addition, there was one DEG with a negative R^2^ change for the 50 μg/kg Mixed BP exposure group and licking/grooming compared to the Corn Oil group, which included Hmga1-ps2 (Supplementary Figure 8B; FDR < 0.05).

Count data from 36,350 genes were used to construct networks for the WCGNA and were clustered into 36 eigengene modules (Supplementary Figure 9). A total of eight eigengene modules were examined for differences between prenatal treatments and postnatal maternal care and their interactions. There were significant main effects or interactions found with two eigengene modules (Black and Yellow).

For the black module (containing 1,097 genes), there was a significant interaction between the 150 μg/kg Mixed BP exposure group and licking/grooming (t = 2.183, p = 0.0411) compared to the Corn Oil group (Figure 8A). The top GO terms for the genes in the black module were Translation, Macromolecule Biosynthetic Process, Peptide Biosynthetic Process, Cytoplasmic Translation, Proteasomal Protein Catabolic Process, and Ubiquitin-Dependent Protein Catabolic Process. The top enriched transcription factors involved in protein-protein interactions with genes in the black module were ESR1, ILF3, POLR2A, MYC, and FOXP3 (Figure 8A). The top enriched transcription factors involved in facilitating transcription of genes in the black module were TAF1, BRCA1, GABPA, CREB1, and YY1 (Figure 8A).

For the yellow module (containing 1,658 genes), there was a significant difference between the 150 μg/kg Mixed BP exposure group and Corn Oil group (t = 2.284, p = 0.0335) and a marginal difference between the 50 μg/kg Mixed BP exposure group and Corn Oil group (t = 1.923, p = 0.0688; Figure 8B). There was also a significant interaction between the 150 μg/kg Mixed BP exposure group and licking/grooming (t = 2.205, p = 0.0393) compared to the Corn Oil group (Figure 8B). The top GO terms for the genes in the yellow module were Translation, Macromolecule Biosynthetic Process, Gene Expression, and Mitochondrial ATP Synthesis Coupled Electron Transport. The top enriched transcription factors involved in protein-protein interactions with genes in the yellow module were ILF3, ESR1, MYC, ILF2, and ESR2 (Figure 8B). The top enriched transcription factors involved in facilitating transcription of genes in the yellow module were TAF1, NFYB, YY1, GABPA, and ELF1 (Figure 8B).
